# Dysregulation of N-terminal acetylation causes cardiac arrhythmia and cardiomyopathy

**DOI:** 10.1101/2023.07.02.546740

**Authors:** Daisuke Yoshinaga, Rui Feng, Maksymilian Prondzynski, Kevin Shani, Yashasvi Tharani, Joseph Milosh, David Walker, Chrystalle Katte Carreon, Bridget Boss, Sheila Upton, Kevin Kit Parker, William T. Pu, Vassilios J. Bezzerides

**Affiliations:** Department of Cardiology, Boston Children’s Hospital, Harvard Medical School, Boston, MA, USA; Disease Biophysics Group, Wyss Institute for Biologically Inspired Engineering, Harvard John A. Paulson School of Engineering and Applied Sciences, Allston, MA USA; Department of Pathology, Boston Children’s Hospital, Harvard Medical School, Boston, MA, USA; Department of Pediatric Cardiology, Dartmouth Hitchcock Medical Center, Manchester, NH USA; Department of Medical Genetics, Dartmouth Hitchcock Medical Center, Manchester, NH USA

## Abstract

**BACKGROUND:** N-terminal-acetyltransferases catalyze N-terminal acetylation (Nt-acetylation), an evolutionarily conserved co-translational modification. Nt-acetylation regulates diverse signaling pathways, yet little is known about its effects in the heart. To gain insights, we studied NAA10-related syndrome, in which mutations in NAA10, which catalyzes Nt-acetylation, causes severe QT prolongation, hypotonia, and neurodevelopmental delay.

**METHODS:** We identified a missense variant in NAA10 (c.10C>A; p.R4S) that segregated with severe QT prolongation, arrhythmia, cardiomyopathy, and sudden death in a large kindred. We developed patient-derived and genome-edited human induced pluripotent stem cell (iPSC) models and deeply phenotyped iPSC-derived cardiomyocytes (iPSC-CMs) to dissect the mechanisms underlying NAA10-mediated cardiomyocyte dysfunction.

**RESULTS:** The NAA10-R4S mutation reduced enzymatic activity, decreased expression levels of NAA10/NAA15 proteins, and destabilized the NatA complex. In iPSC-CM models of NAA10 dysfunction, dysregulation of the late sodium and slow rectifying potassium currents caused severe repolarization abnormalities, consistent with clinical QT prolongation and increased risk for arrhythmogenesis. Engineered heart tissues generated from mutant NAA10 cell lines had significantly decreased contractile force and sarcomeric disorganization, consistent with the cardiomyopathic phenotype in the identified family members. Diastolic calcium levels were increased with corresponding alterations in calcium handling pathways. We identified small molecule and genetic therapies that reversed the effects of NAA10 dysregulation of iPSC-CMs.

**CONCLUSIONS:** Our study defines novel roles of Nt-acetylation in cardiac ion channel regulation and delineates mechanisms underlying QT prolongation, arrhythmia, and cardiomyopathy caused by NAA10 dysfunction.

## INTRODUCTION

N-terminal-acetyltransferases (NATs) catalyze protein N-terminal acetylation (Nt-acetylation), an evolutionarily conserved co-translational modification that regulates protein degradation, protein-protein interactions, membrane targeting, and protein folding^1^. After the initiation of translation, methionine aminopeptidases excise the initiator methionine from 80% of mammalian proteins, creating a substrate for NATs to irreversibly transfer an acetyl group from acetyl-CoA. N-terminal-acetyltransferase A (NatA), one of five mammalian NATs with distinct substrate specificities^2^, preferentially acetylate G, S, A, T, or V residues exposed at protein N-termini by α-methionine removal^3^. By sequence analysis, NatA can modify up to 40% of expressed mammalian proteins, although there are few functionally validated targets^4^. Despite the diversity of affected signaling pathways, little is known about the effects of Nt-acetylation in the heart.

NatA contains a catalytic subunit, NAA10, and a regulatory subunit, NAA15. Pathological variants in NAA10 cause NAA10-related syndrome, a rare multi-system disorder characterized by developmental delay, hypotonia, QT prolongation, arrhythmias, and increased mortality often before 1 year of age^5^. In addition to QT prolongation and sudden death, NAA10 variants are associated with hypertrophic cardiomyopathy and congenital heart defects including atrial septal defects, ventricular septal defects, and tetralogy of Fallot^5–13^. Located on the X-chromosome, pathogenic NAA10 variants are inherited in an X-linked recessive pattern, with severe clinical phenotypes manifesting predominantly in males^6, 8, 9, 12, 13^. Female carriers present with more variable manifestations that range from mild to severe developmental delay and cardiac involvement^8, 10, 12^. Patients with cardiac involvement more commonly have NAA10 variants within the N-terminally located NAA15 interaction or catalytic domains^5, 7–9^, whereas those with neurodevelopmental delays generally have variants located closer to the C-terminus^14^, suggesting the importance of NatA complex formation for cardiac homeostasis.

Recent large-scale exome sequencing projects of patients with congenital heart disease identified several *de novo* NAA15 variants^15^. Cardiomyocytes derived from human induced pluripotent stem cells (iPSC-CMs) with a single missense mutation (NAA15^R276W/WT^) or NAA15 haploinsufficiency (NAA15^WT/-^) demonstrated minor contractile defects under loaded conditions, whereas iPSCs lacking NAA15 had poor viability and did not readily differentiate into iPSC-CMs^15^. These data support the importance of NatA function in a broad range of cardiovascular diseases, but the underlying mechanisms and cardiac targets remained undefined.

Despite NAA10-related syndrome pointing to a key role of NAA10 and Nt-acetylation in human cardiac homeostasis, identification of cardiac-specific NatA protein targets has been limited. Here using iPSC disease-modeling, bioengineering, cellular electrophysiology, and optogenetics; we investigate the mechanisms by which an NAA10 missense variant causes long QT syndrome and cardiomyopathy. We identify the first cardiac-specific targets dysregulated by NAA10 dysfunction and provide novel mechanistic insights into the roles of Nt-acetylation in cardiac homeostasis and disease.

## METHODS

The extended Methods section in the online-only Data Supplement provides additional information. All data and materials that support the findings of this study are available from the corresponding author on reasonable request.

### Patient Data

The proband was identified as part of a large family originally diagnosed with “gene-negative” long QT syndrome as a referral to the clinical practice of VJB. Retrospective clinical data was collected after enrollment into an Institutional Review Board (IRB) approved protocol at Boston Children’s Hospital. A four-generation family history was performed by a certified genetic counselor (S.U.) and all clinical information was collected and collated by a registered nurse study coordinator (B.B.). Echocardiography, electrocardiography, and remote device monitoring were performed as standard of care at Boston Children’s Hospital and Dartmouth Hitchcock Medical Center. Postmortem examination with cardiac histopathological evaluation was performed by the Cardiac Registry service (C.K.C.) at Boston Children’s Hospital.

### Generation of iPSC lines

Patients from a clinical cohort of NAA10-related syndrome consented to participate in this study, supplied peripheral blood mononuclear cells (PBMCs) for somatic-cell reprogramming into iPSCs. The introduction of the NAA10^R4S^ variant through genome-editing in a control background (WTC11) generated an isogenic iPSC model. Variant sequencing, pluripotency marker staining, and karyotype analysis performed at regular intervals ensured iPSC model integrity.

### Electrophysiology

To measure the electrical effects of iPSC monolayers, we plated iPSC-CMs on 24 well multi-electrode array (MEA) plates (Axion Biosystems). After 48 hours in culture, adenoviral transduction for the channelrhodopsin ChR2 fused to green fluorescent protein (Ad-ChR2-GFP) enabled optical pacing. We performed optical pacing at 1Hz using a multi-well array LED and measured the resulting extracellular field potential (FP). The time from the onset of the FP to the peak of the repolarization defined the field potential duration (FPD).

For single cell electrophysiology, isolated iPSC-CMs were plated on coated coverslips. Borosilicate glass pipettes pulled to form recording electrodes with a resistance of 2 to 3 MΩ were used to establish whole-cell recordings either with direct membrane rupture or with the inclusion of β-escin (25μM) in the recording solution to create the perforated patch configuration. Selective channel blockers and stereotypical voltage protocols isolated individual currents (Supplemental table 1). All the data were acquired from at least three independent experiments using different biological replicates.

### Structural analyses and bioengineering

To quantify structural effects of NAA10 dysfunction, we performed a hierarchal examination of cardiomyocyte structure and contractile function. For single-cell analysis we plate dissociated iPSC-CMs on PDMS-coated glass coverslips with micro-contacted printed islands of extra-cellular matrix (ECM) with a 7:1 aspect ratio. After fixation, cells were stained for sarcomeric alpha actinin and actin for confocal imaging and automated structural analysis.

To measure contractile defects, 3D heart tissue constructs formed from iPSC-CMs in a fibrin gel and molded between silicone pillars enabled the assessment of contractile force. A custom-built microscope imaged engineered heart tissues at video-rates for subsequent automated motion analysis. Periodic optical stimulation at 1Hz with 488 nm light after transduction with adenovirus for ChR2-GFP, normalized EHT contraction rates at 1Hz for group statistics.

## RESULTS

### Identification of novel NAA10 variant

We evaluated a four-generation kindred with multiple members who had QT prolongation and sudden cardiac death (SCD) in young patients (Figure 1A). Genetic testing for mutations associated with congenital long QT syndrome (cLQTS) revealed a variant of unknown significance in KCHN2 p.R164H (Clinvar, VCV000067508.8) that did not segregate with the disease phenotype (Figure 1A). In addition to QT prolongation and arrhythmia, male patients had significant neurodevelopmental delay and mild peripheral myopathy. These extra-cardiac phenotypes suggested a multi-system disorder. Expanded DNA sequencing identified a novel variant, *NAA10* c.10 C>A, p.R4S, that co-segregated with clinical manifestations in an X-linked recessive pattern in family members available for genetic testing (Figure 1B). Male family members with the *NAA10^R4S^* variant had significant QT prolongation and T-wave abnormalities (Patient III-10, Figure 1C). The NAA10^R4S^ variant lies within the N-terminal NAA15 interaction domain necessary for formation of the NatA complex^16^. Consistent with severe QT prolongation, patient III-10 developed frequent episodes of ventricular tachycardia despite high-dose beta-blocker therapy (nadolol 80 mg twice daily), necessitating insertion of an implantable cardiac defibrillator (ICD) (Figure 1E). All his episodes of ventricular arrhythmia occurred during rest or with only minimal physical activity. This patient developed progressive heart failure with an ejection fraction < 25%, LV dilation, and symptoms consistent with class IV NYHA heart failure^17^. Despite aggressive medical and supportive therapies, he died from complications of heart failure at 17 years of age (Figure 1F). Histopathology of the post-mortem LV samples demonstrated prominent interstitial fibrosis, and hypertrophic cardiomyocytes with bizarre hyperchromatic nuclei and highly irregular nuclear contours (Figure 1G). These clinical data support the identification of a novel NAA10 variant associated with a severe cardiac disease phenotype.

**Figure 1.**
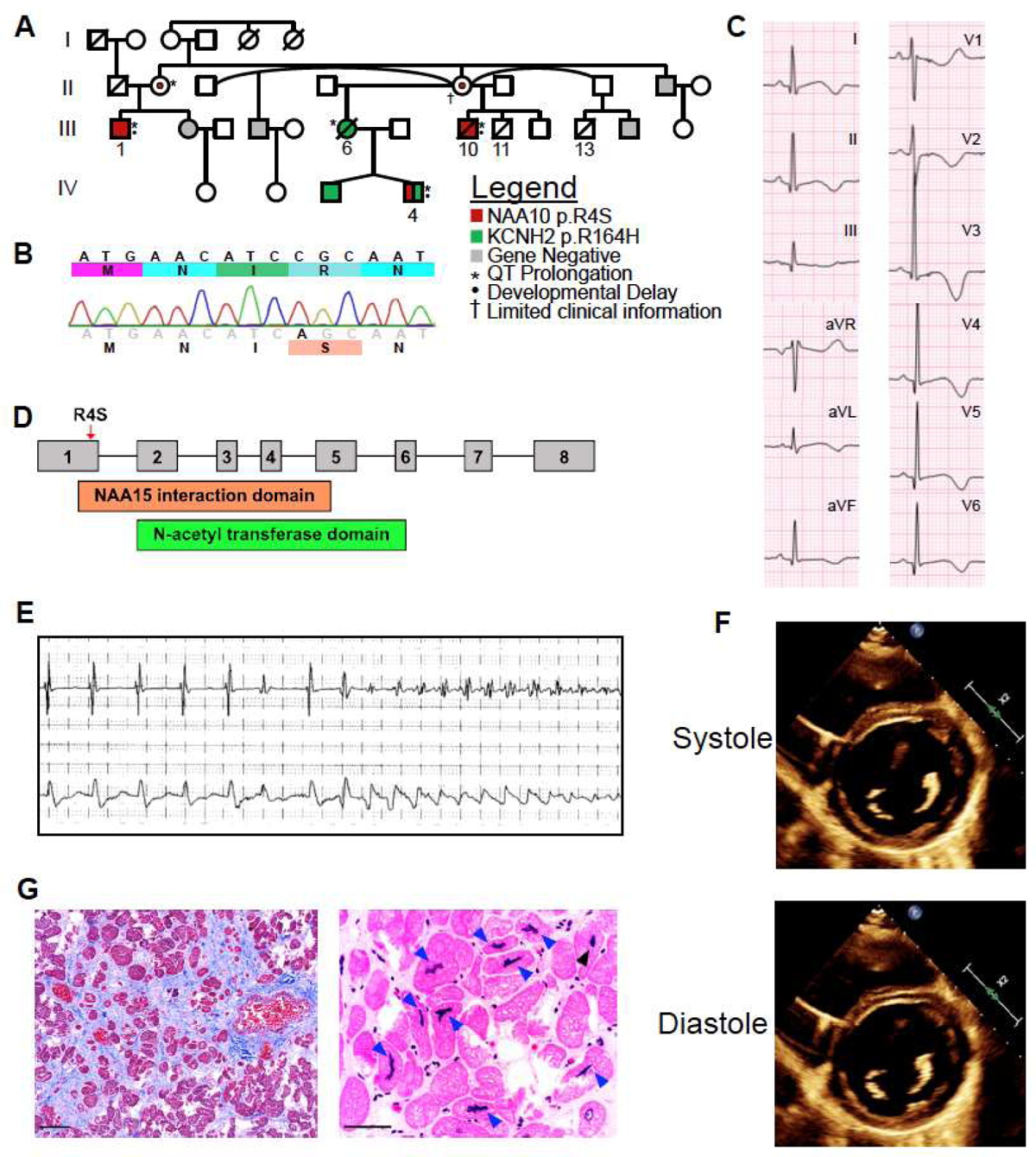
A novel NAA10 variant is associated with clinical long QT syndrome and severe cardiomyopathy. **A.** A four generation family pedigree identified predominately male patients with QT prolongation, developmental delay, and early mortality in a family referred for “gene-negative long QT syndrome”. Clinical sequencing revealed the NAA10 p.R4S variant (red) to segregate with the clinical phenotype, whereas the variant of unknown significance in KCNH2 (p.R164H, shown in green) did not segregate with QT prolongation. In generation III: both male subjects (III-11, III-13) died as infants. Female (III-6) died at the age of 26 years from cardiac arrest prior to the availability of testing for NAA10. **B.** Sanger sequencing of the proband (indicated by arrow). **C.** ECG for patient III-10 with QT prolongation (QTc = 545 msec) and inverted T-waves. **D.** A schematic of the exon structure of the NAA10 gene overlaid with the novel p.R4S variant and its relationship to the domain structure of the NAA10 protein. **E.** Tracing from implantable cardiac defibrillator with short-long-short coupling prior to initiation of Torsades de Pointes (TdP). **F.** Short axis echocardiography of patient III-10. Calculated ejection fraction (EF) of 22.3%. **G.** Cardiac micrographs from patient III-10 post-mortem. Right panel shows Masson Trichrome staining (scale bar = 100 µm) and left panel is hematoxylin and eosin staining (scale bar = 50 µm) of left ventricle. Arrows indicate abnormal appearing nuclei.

### The R4S mutation impairs NatA catalytic activity

The arginine at position 4 (R4) is highly conserved across different species, suggesting a key role in NAA10 function (Figure 2A). *In silico* analysis of existing high-resolution structures of the NAA10-NAA15 complex^18^ predicted that the R4S mutation would impair local hydrogen bonding and ionic interactions and destabilize NAA10 (Figure 2B). In addition to effects on NAA10 stability, the R4S variant could also interfere with the interaction surface between NAA10 and NAA15 (Figure 2B). Because of the predicted disruption of the NAA10/NAA15 interaction domain, we investigated NatA complex formation. In transfected HEK293 cells, NAA10-R4S co-immunoprecipitated 50% less NAA15 than wild-type NAA10 (Figure 2C). Furthermore, we noted that NAA10-R4S expressed less protein than wild-type (Supplemental Figure 1A), leading us to suspect increased protein degradation. To assess protein half-life, we inhibited protein synthesis with cycloheximide and measured protein levels of expressed WT and mutant NAA10 proteins^8^. NAA10-R4S displayed more rapid protein degradation than NAA10-WT when normalized to vinculin (Figures 2D and 2E).

**Figure 2.**
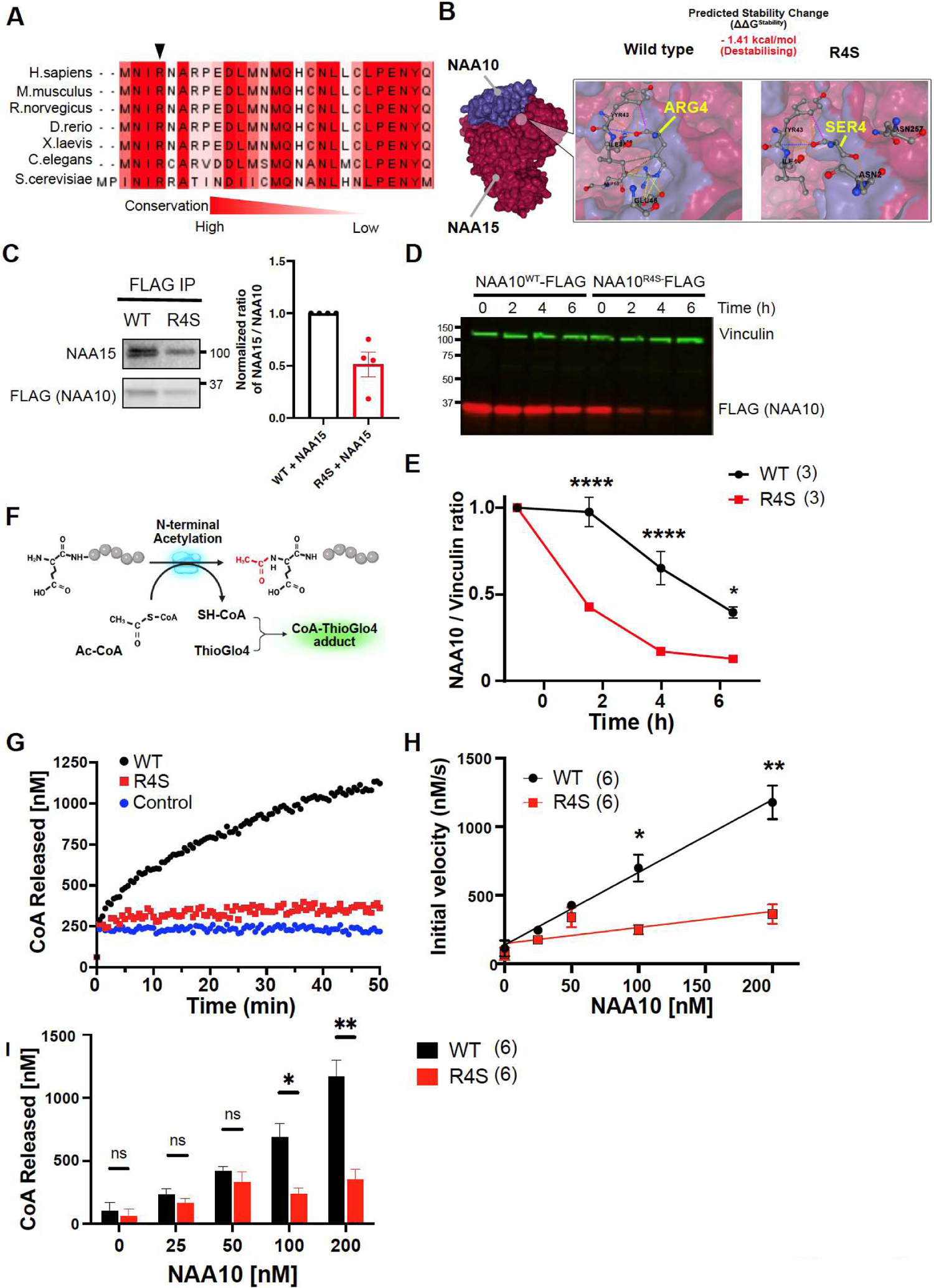
The R4S mutation induces NAA10 instability and decreases enzymatic function. **A.** Alignment of NAA10 protein sequences across multiple species. The arginine indicated by the arrow is highly evolutionarily conserved. **B.** HEK293 cells were co-transfected with NAA10-FLAG and NAA15 and lysates were immunoprecipitated with anti-FLAG antibody. Immunoprecipitants were separated by gel electrophoresis with subsequent immunoblotting using anti-NAA10 and anti-NAA15 antibodies. The binding capacity is quantified as a function of NAA10 protein levels (right panel). **C.** The R4S mutation alters NAA10 protein stability and possible interactions with NAA15 by *in silico* predication (DynaMut2). **D.** HEK293T cells transfected with NAA10^WT^-FLAG and NAA10^R4S^-FLAG were treated with cycloheximide for 0, 2, 4, and 6 hours and cell lysates were analyzed by western blotting with fluorescent secondary antibodies for vinculin and FLAG. **E.** Quantification of D, with normalization of NAA10/vinculin ratio to initial NAA10 protein levels. **F.** Schematic of ThioGlo4 assay to quantify the enzymatic activity of NAA10. Enzymatic production of CoA is detected by ThioGlo4 probe with excitation at 400 nm and emission at 465 nm. **G.** Representative traces of ThioGlo4 reaction over time indicating NAA10 activity. NAA10 WT or NAA10-R4S enzyme concentrations were 100nM. No enzyme was added to the control sample. **H.** The initial velocity of the enzymatic reaction was measured ThiolGlo4 fluorescence within the first 10 seconds. **I.** The total amount of CoA released was measured at 30 minutes after initiation of the enzymatic reaction. Statistics performed by one-ANOVA with Tukey’s post-hoc test for B. ****P <0.0001, **P<0.01, *P<0.05.

As the catalytic subunit of NatA, NAA10 catalyzes the transfer of acetyl-CoA to protein N-termini, releasing free CoA^12^. Free CoA reacts with ThioGlo4, forming a fluorescent adduct (Figure 2F). We used this reaction to measure the effect of the R4S mutation on NAA10 catalytic activity. Monitoring ThioGlo4-CoA fluorescence over time showed that purified NAA10-R4S protein had significantly lower catalytic activity than NAA10-WT protein (Figures 4G and 4H). Increasing protein concentration failed to overcome the enzymatic defect of NAA10-R4S even at the termination of the reaction (Figure 2I). Collectively these data show that the novel mutation NAA10-R4S destabilizes the NatA complex, increases NAA10 protein degradation, and directly impairs enzymatic activity.

### Increased risk of arrhythmogenesis in a model of NAA10 dysfunction

To model the effects of NAA10-R4S and gain mechanistic insight into NAA10 function in cardiomyocytes, we created an induced pluripotent stem cell lines (iPSC) from the patients III-1 and III-10 through somatic-cell reprogramming^19^. Isolated iPSC clones were positive for the NAA10^R4S^ variant and had normal karyotypes and markers of pluripotency (Supplemental Figures 2A-2C). A single patient-derived clone from patient III-10 was selected for further studies and was designated as pNAA10^R4S^. To control for genetic background, we also introduced the NAA10^R4S^ variant into a male WT iPSC line by genome editing. This genome-edited isogenic line was designated as eNAA10^R4S^ (Supplemental Figure 2D). We also sequenced gene predicted to be off-target sites for genome editing and did not detect any additional mutations (Supplemental Figure 2E). We differentiated NAA10 mutant and WT iPSC-lines into iPSC-CMs using established small molecule differentiation protocols^20^. We confirmed robust cardiomyocyte differentiation of all three iPSC lines by cardiac troponin T (cTnT) immunostaining followed by flow cytometry (Supplemental Figure 2F). Western blotting of both pNAA10^R4S^ and eNAA10^R4S^ iPSC-CMs showed that the mutant lines expressed 50% less NAA10 protein compared to the control line (Figures 3A and 3B), consistent with decreased stability of NAA10-R4S.

**Figure 3.**
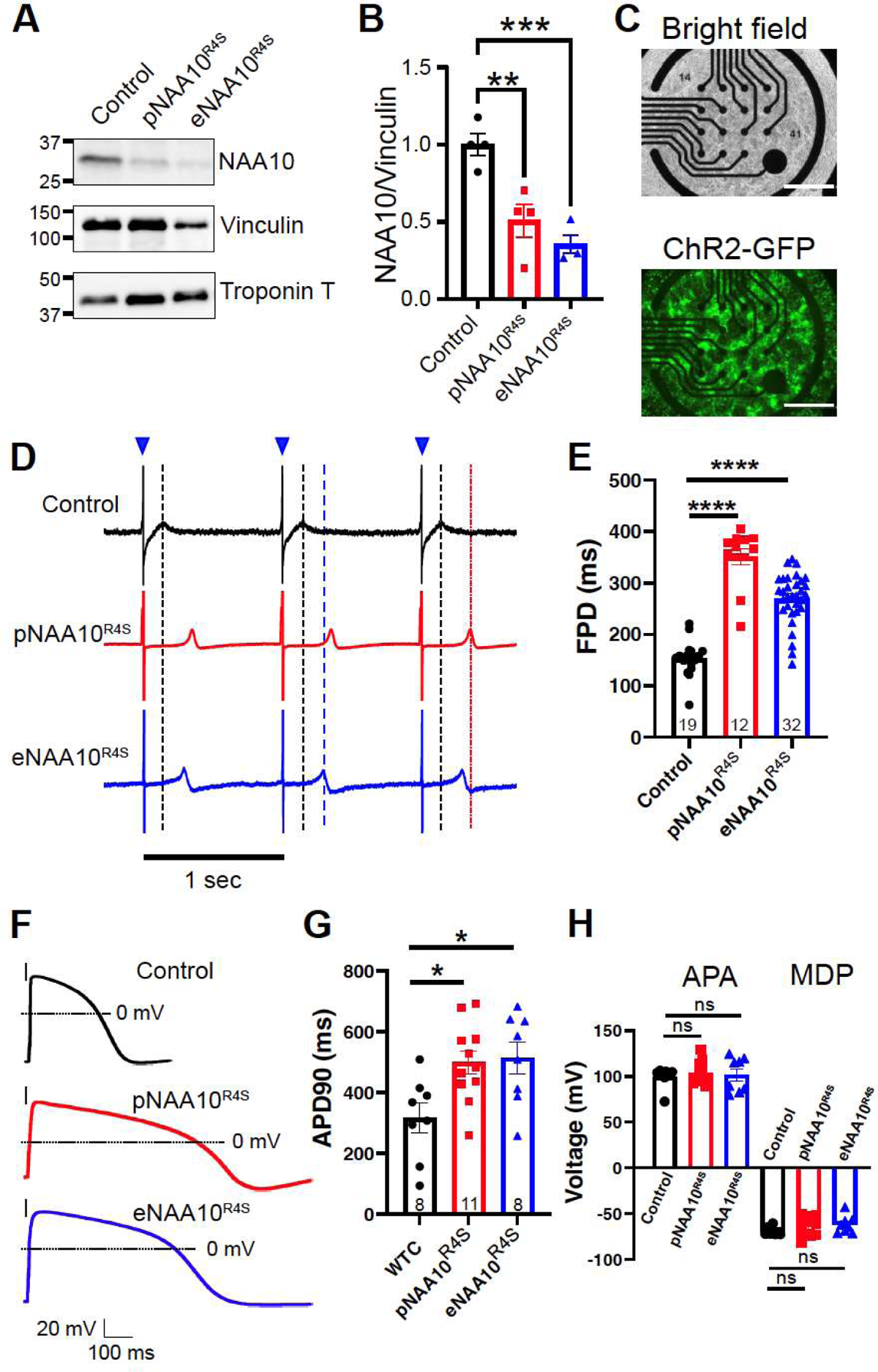
A cellular model of NAA10 dysfunction demonstrates repolarization abnormalities. **A.** Whole cell lysates from differentiated control, pNAA10^R4S^, or eNAA10^R4S^ iPSC-CMs were separated by gel electrophoresis and subjected to Western blot analysis with antibodies specific to native NAA10, vinculin or cardiac troponin T. **B.** Quantification of total NAA10 protein levels were normalized to vinculin. **C.** Differentiated iPSC-CMs were plated on multi-electrode arrays (MEAs, upper panel) and transduced with adenovirus for the channelrhodopsin ChR2-GFP (lower panel) for optical pacing. Scale bar = 750μm **D.** Representative MEA recordings of control iPSC-CMs or harboring the NAA10 pR4S variant as either patient-derived (pNAA10^R4S^) or genome-edited (eNAA10^R4S^) iPSC-CMs with optical pacing at 1Hz with 488nm light (blue triangles). Dashed lines represent the field potential duration (FPD) for control (black), pNAA10^R4S^ (red), and eNAA10^R4S^ (blue) iPSC-CMs. **E.** Quantification of FPD paced at 1Hz from B. FPD of pNAA10^R4S^-, and eNAA10^R4S^-iPSC-CMs. **F.** Representative whole-cell recordings of single iPSC-CMs under current clamp conditions, electrically paced at 1Hz. **G.** Quantification of the action potential duration (APD) at 90% of the peak membrane potential demonstrates significant prolongation in both pNAA10^R4S^ and eNAA10^R4S^ iPSC-CMs compared to control cells. **H.** The peak action potential amplitude (APA) and mean diastolic potential (MDP) were not affected by the NAA10^R4S^ variant. The number of analyzed samples (n) is annotated on each graph (sample numbers on panel G are the same for H), which were from at least three separate differentiations. Statistics were performed by the one-way Kruskal-Wallis test with Dunnet’s test for multiple comparisons: ns=p>0.1, *p<0.05, **p<0.01, ***p<0.005, ****p<0.0001.

As previously described, NAA10-related syndrome is associated with QT prolongation and SCD, but the underlying mechanisms are poorly understood. To determine if our iPSC models recapitulated the observed clinical phenotypes, we plated WT, pNAA10^R4S^ and eNAA10^R4S^ iPSC-CMs on multi-electrode arrays (MEAs). The field potential duration (FPD) measured from iPSC-CMs monolayers correlates with the EKG-based QT interval^21^. Since FPD depends on iPSC-CM beat rates^22^, we used optical pacing to pace iPSC-CMs at a fixed rate. We transduced iPSC-CMs with adenovirus expressing the channelrhodopsin ChR2 fused to GFP (Ad-ChR2-GFP, Figure 3C) and optically paced monolayers with 488nm light at 1 Hz (Figure 3D, blue triangles). Under these conditions, pNAA10^R4S^ and eNAA10^R4S^ iPSC-CMs had dramatically prolonged FPDs compared to WT cells (WT = 150.5 ± 7.9 msec; pNAA10R4s = 349.7 ± 17.4 msec; eNAA10R4S = 268.3 ± 9.2 msec; P<0.0001; Figures 3D and 3E).

FPD and QT interval are each measures of cardiomyocyte repolarization. To evaluate the effect of NAA10-R4S more directly on cardiomyocyte repolarization, we performed single-cell electrophysiology experiments on iPSC-CMs. After establishing a whole-cell current clamp configuration, we injected depolarizing currents at 1 Hz to elicit membrane action potentials (APs). iPSC-CM evoked APs had a characteristic ventricular-like morphology. The action potential duration (APD) measured at 90% of peak repolarization (APD90) was more than 2-fold longer in NAA10-mutant compared to WT iPSC-CMs (Figures 3F and 3G). There were no differences in action potential amplitude (APA) or mean diastolic potential (MDP) between WT and NAA10^R4S^ mutant iPSC-CMs, suggesting the observed APD prolongation did not arise from differences in differentiation or maturation^23^. Collectively these data demonstrate that the NAA10^R4S^ variant causes severe repolarization abnormalities in iPSC-CMs.

### NAA10 dysfunction dysregulates both sodium and potassium currents

Congenital long QT syndrome (cLQTS) is characterized QT prolongation on the surface EKG and increased arrhythmogenic risk^24^. Underlying genetic variants within ion channel or ion channel-related proteins are identified in up to 80% of cLQTS patients^25^. To identify potential target proteins that might be affected by NAA10 dysregulation and cause repolarization abnormalities, we exploited the discovery that N-terminal protein sequence predicts the likelihood of Nt-acetylation. Therefore, we analyzed the N-terminal sequences of all proteins associated with cLQTS and their respective probability of modification by NatA (Table 1). These data suggested that multiple cardiac ion channels and related proteins known to underlie the cardiac action potential, could be NatA targets and therefore affected by NAA10 dysfunction.

**Table 1.**
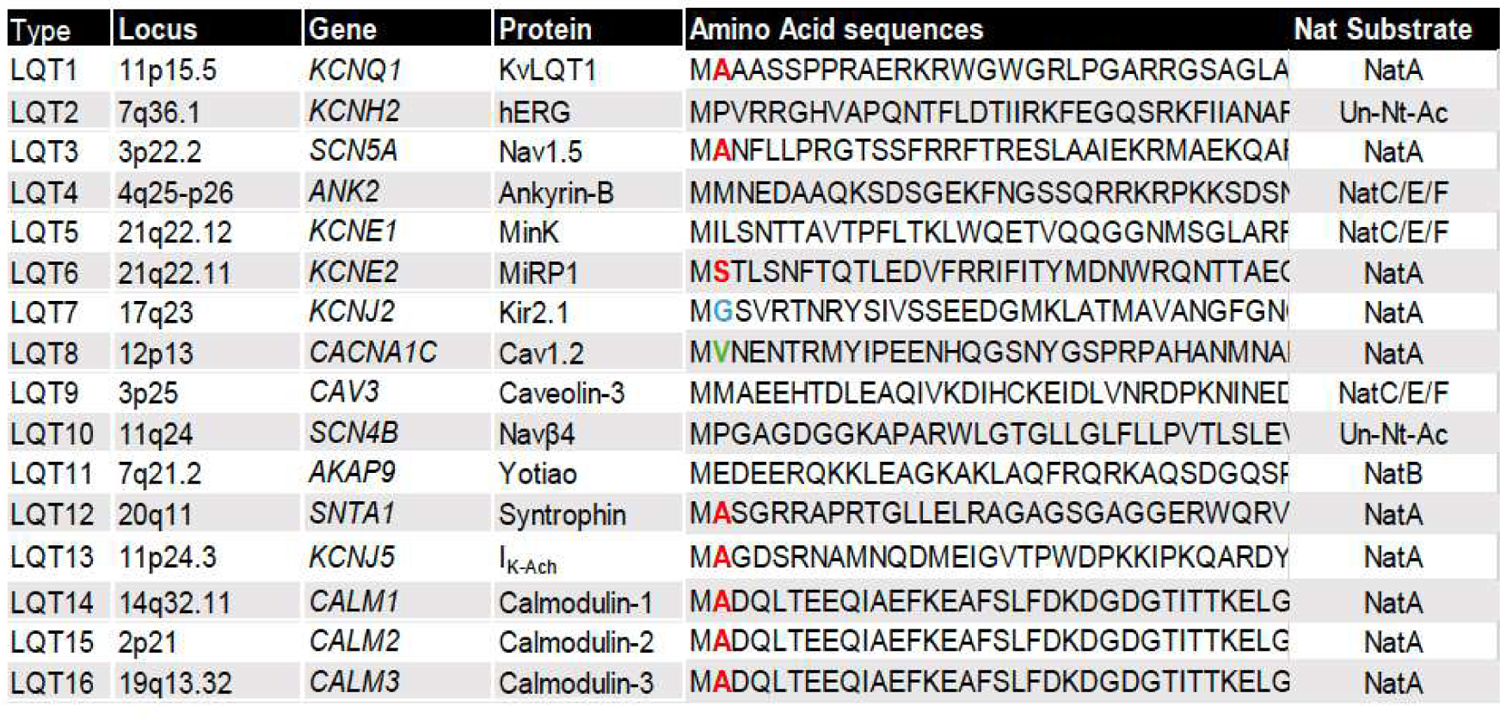
Amino and sequences for the LQT-related poroteins

We systematically investigated the major currents responsible for the cardiac action potential. The voltage-gated sodium channel SCN5A (protein: Na_V_1.5), underlies the rapid depolarization of the cardiac action potential. SCN5A gain of function variants cause LQT type III (LQT3)^26^ through at least two mechanisms: (1) alterations in voltage dependent activation/inactivation and (2) increased late sodium current.

To investigate the first mechanism, in the presence of the calcium channel blocker nifedipine (10 μM), depolarizing steps from a hyperpolarized holding potential of −100mV in single iPSC-CMs evoked stereotypical rapidly activating and deactivating currents consistent with I_Na_ (Figure 4A). Peak inward sodium currents were significantly increased in pNAA10^R4S^ and eNAA10^R4S^ iPSC-CMs by as much as 2.5-fold compared to WT cells (Figure 4A and 4B upper panel). While increased I_Na_ current density is associated with heart failure and other forms of Na_V_1.5-mediated cardiovascular disorders, it is not a recognized mechanism for cLQTS. Instead, differential alterations in the voltage-dependent activation and inactivation of I_Na_ can increase the “window current” and are associated with LQT3. While there was a small shift to more hyperpolarizing potentials in both the activation and inactivation curves in NAA10-mutant iPSC-CMs there was no increase in the net activation probability (Figure 4B lower panel, Supplemental Table 2).

**Figure 4.**
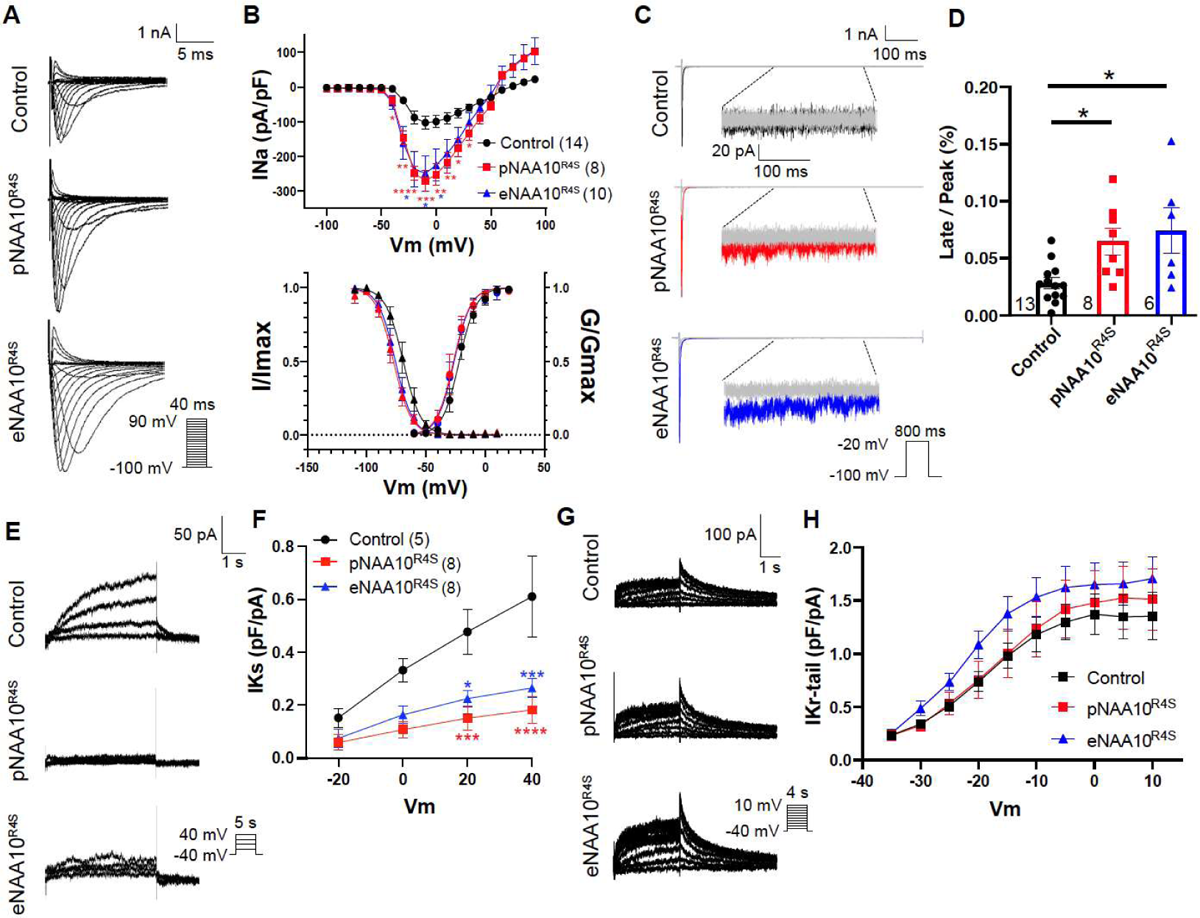
Increased late I_Na_ and decreased I_Ks_ contributes to action potential prolongation in NAA10-mutant iPSC-CMs. **A.** Representative traces of sodium current (I_Na_) under whole-cell voltage-clamp conditions from control, pNAA10^R4S^, and eNAA10^R4S^-iPSC-CMs. **B.** Current-voltage relation of normalized whole-cell I_Na_ with peak current densities significantly increased in both pNAA10^R4S^, and eNAA10^R4S^-iPSC-CMs (upper panel). I_Na_ normalized with maximal conductance and maximal current density represent voltage-dependent activation and inactivation curves respectively (lower panel). **C.** Representative whole-cell current traces in response to long depolarizing pulses to elicit the persistent or late sodium current (I_NaL_) and normalized to tetrodotoxin blockade (TTX, gray traces). **D.** Quantification of I_NaL_ normalized to peak sodium current (I_NaP_) after leak subtraction with TTX. **E.** Representative whole-cell traces of the slow-rectifying potassium current (I_Ks_) from control and mutant NAA10 iPSC-CMs. **F.** The I_Ks_ current-voltage relationship demonstrated decreased peak outward current densities in pNAA10^R4S^, and eNAA10^R4S^-iPSC-CMs as compared to control cells. **G.** Representative whole-cell current traces of the rapid-activating potassium current (I_Kr_) in response to increasing voltage steps. **H.** After channel activation, the whole-cell voltage was stepped to the same voltage and the measured Instantaneous current was quantified as the I_Kr_-tail current for control, pNAA10^R4S^, and eNAA10^R4S^ iPSC-CMs. The number of cells (n) is annotated on each graph and samples were from at least three independent differentiations. Statistics were performed by one-way Kruskal-Wallis with Dunnett’s multiple comparisons test: ns=p>0.1, *p<0.05, **p<0.01, ***p<0.005, ****p<0.0001.

The second most common mechanism for LQT3 is the increase in persistent or late sodium current (I_NaL_) which has also been associated with heart failure^27^. To investigate the effects of the NAA10^R4S^ variant and associated NAA10 dysfunction on I_NaL_, we stimulated iPSC-CMs with long-depolarizing steps (1s) at baseline and in the presence of 30 µM tetrodotoxin (TTX) to normalize for background membrane leak. Consistent with a LQT3-type phenotype there was a >2-fold increase in I_NaL_ in pNAA10^R4S^ and eNAA10^R4S^ iPSC-CMs as compared to WT iPSC-CMs (Figures 4C and 4D) suggesting a direct biophysical effect of NAA10 on Na_V_1.5 channel function.

cLQTS is also caused by alterations in potassium flux, which drives cardiomyocyte repolarization^25^. To investigate dysregulation of potassium currents as contributing to APD prolongation in NAA10-related syndrome, we performed voltage-clamp whole-cell recordings of single iPSC-CMs and isolated the two major repolarizing potassium currents, slow activating (I_Ks_) and rapid activating (I_Kr_). We isolated I_Ks_ by including HMR1556 (1 µM) in the recording solution to block KCNH2 channels (protein, K_V_11.1; Figure 4E). Activating voltage steps demonstrated a significant reduction in I_Ks_ current density in eNAA10^R4S^ and pNAA10^R4S^ iPSC-CMs within the physiologic depolarization range (Figures 4E and 4F). Next, we selectively isolated I_Kr_ by incubation with E403 (1 µM) to block KCNQ1 channels (protein K_V_7.1). This maneuver did not reveal significant differences I_Kr_ in NAA10^R4S^ mutant compared to WT iPSC-CMs (Figures 4G, 4H and supplemental Figure 3A).

Alterations in the inward calcium (Ca^2+^) current I_Ca-L_ also cause cLQTS (Timothy Syndrome, LQT8)^28^. We recorded L-type Ca^2+^ channels by inhibiting Na_V_1.5 channels through high-dose TTX in the recording bath and an elevated holding potential (−40mV). We did not observe significant differences in total current density or L-type Ca^2+^ channel properties in NAA10-mutant iPSC-CMs (Supplemental Figures 3C and 3D, Supplemental Table 1). Taken together, these data indicate that QT prolongation and risk for arrhythmia in NAA10-related syndrome are caused by a combination of increased I_NaL_ and reduced I_Ks_.

### NAA10^R4S^ iPSC-CMs have structural and contractile abnormalities

Patients with NAA10-related syndrome have variable heart disease phenotypes ranging from cardiomyopathy to structural heart disease^46^. Our patient III-10 died from complications related to severe dilated cardiomyopathy (DCM) which has not been previously associated with NAA10-related syndrome. To investigate possible cardiomyopathic effects of the *NAA10^R4S^* variant and related NAA10 dysfunction, we performed detailed imaging analysis of single iPSC-CMs. We used micro-contact printing to deposit rectangular fibronectin islands with the 7:1 aspect ratio characteristic of adult human ventricular cardiomyocytes (Fig 5A)^29–31^. Plating iPSC-CMs on these patterned substrates induces sarcomere alignment, structural integrity, and features of cellular maturity when compared to unpatterned cells (Figure 5B)^31–34^. High-resolution confocal imaging of iPSC-CMs stained for sarcomeric alpha actinin (SAA) and subjected to objective, computational image analysis^35^ demonstrated decreased sarcomeric packing density (SPD) and increased sarcomere length in NAA10^R4S^ mutant iPSC-CMs (Figure 5C and 5D, respectively), consistent with a DCM-like phenotype^36^.

**Figure 5.**
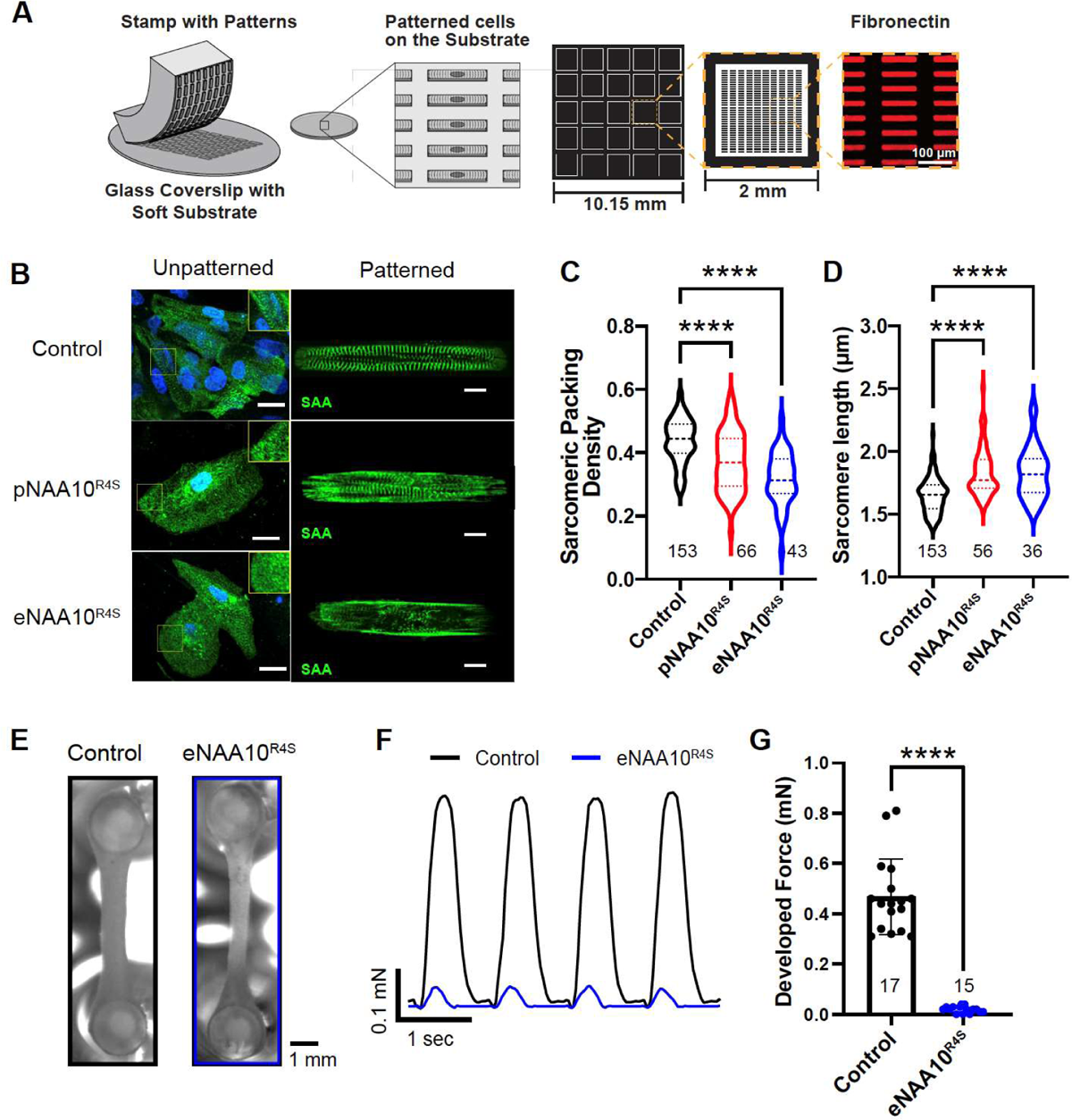
NAA10 mutant iPSC-CMs have severe structural and contractile defects. **A.** Depiction of micro-contact printing with PDMS stamps of fibronectin for spatially restricted patterning of iPSC-CMs. Rectangle patterns have a length to width ratio of 7:1. **B.** Confocal micrographs of Control, pNAA10^R4S^, or eNAA10^R4S^ iPSC-CMs, stained for sarcomeric alpha actinin (SAA) as either unpatterned (left panels, scale bar = 20μm) or patterned on micro-contacted printed substrates (right panels, scale bar = 10μm). **C.** A customized Matlab program calculated **C.** sarcomeric packing density (SPD) and **D.** sarcomere length from single micro-patterned iPSC-CMs. Lower SPD values correspond to a more disordered sarcomeric structure. **E.** Brightfield images of engineered heart tissues (EHTs) created from isogenic control and eNAA10^R4S^ iPSC-CMs on PDMS “pillars” after transduction with ChR2-GFP adenovirus for optical pacing. **F.** Representative traces of developed contractile force of EHTs calculated from video data of PDMS pillar displacement in response to optical pacing with 488nm light at 1Hz. **G.** Quantification of contractile force calculated over 10 beats from two independent differentiations. The number of cells or tissues (n) is annotated on each graph and samples were from at least three independent differentiations unless otherwise specified. Statistics were performed by one-way Kruskal-Wallis with Dunnett’s multiple comparisons test (C and D) or the Mann-Whiney test for G: ns=p>0.1, *p<0.05, **p<0.01, ***p<0.005, ****p<0.0001.

To further investigate the effects of NAA10 dysfunction on contractile force, we generated 3D-engineered heart tissues (EHTs) from WT and eNAA10^R4S^ iPSC-CMs^37^. Cells embedded in extracellular matrix were molded around two silicone pillars and transduced with Ad-ChR2-GFP to facilitate optical pacing (Figure 5E). After tissue formation and maturation, EHTs were optically paced at 1 Hz and imaged. Contractile force was measured based on pillar displacement (Figure 5F). Despite long-term culture for up to 28 days, eNAA10^R4S^ EHTs generated minimal force that was significantly lower than WT EHTs (Figures 5F and 5G). These data collectively identify novel structural defects and a severe contractile defect that was not observed in a previously reported model of NatA dysfunction caused by a NAA15^15^.

### Abnormal Ca^2+^ handling in NAA10^R4S^ iPSC-CMs

Balanced homeostasis of intracellular Ca^2+^ levels is critical for normal cardiomyocyte functioning. Indeed, excess diastolic Ca^2+^ from either decreased cytosolic clearance or increased influx is associated with impaired cardiomyocyte relaxation and contractility^38^. To determine if Ca^2+^ handling defects underlie impaired contractility in NAA10^R4S^ mutant iPSC-CMs, we loaded microcontact-printed iPSC-CMs with the ratiometric calcium indicator Fura-2. With electrical pacing at 0.5 Hz, Ca^2+^ transients in eNAA10^R4S^ iPSC-CMs did not have a significantly different peak height or diastolic Ca^2+^ level (Figures 6A-C). However, with pacing at more a physiologic rate of 1 Hz there was a significant increase in diastolic Ca^2+^ levels (Figures 6A and 6B). Further, the calcium transient relaxation coefficient, a measure of the rate of cytosolic Ca^2+^ clearance, was significantly elevated in the eNAA10^R4S^ iPSC-CMs compared to controls (Figures 6A, 6D, and Supplemental Table 2).

**Figure 6.**
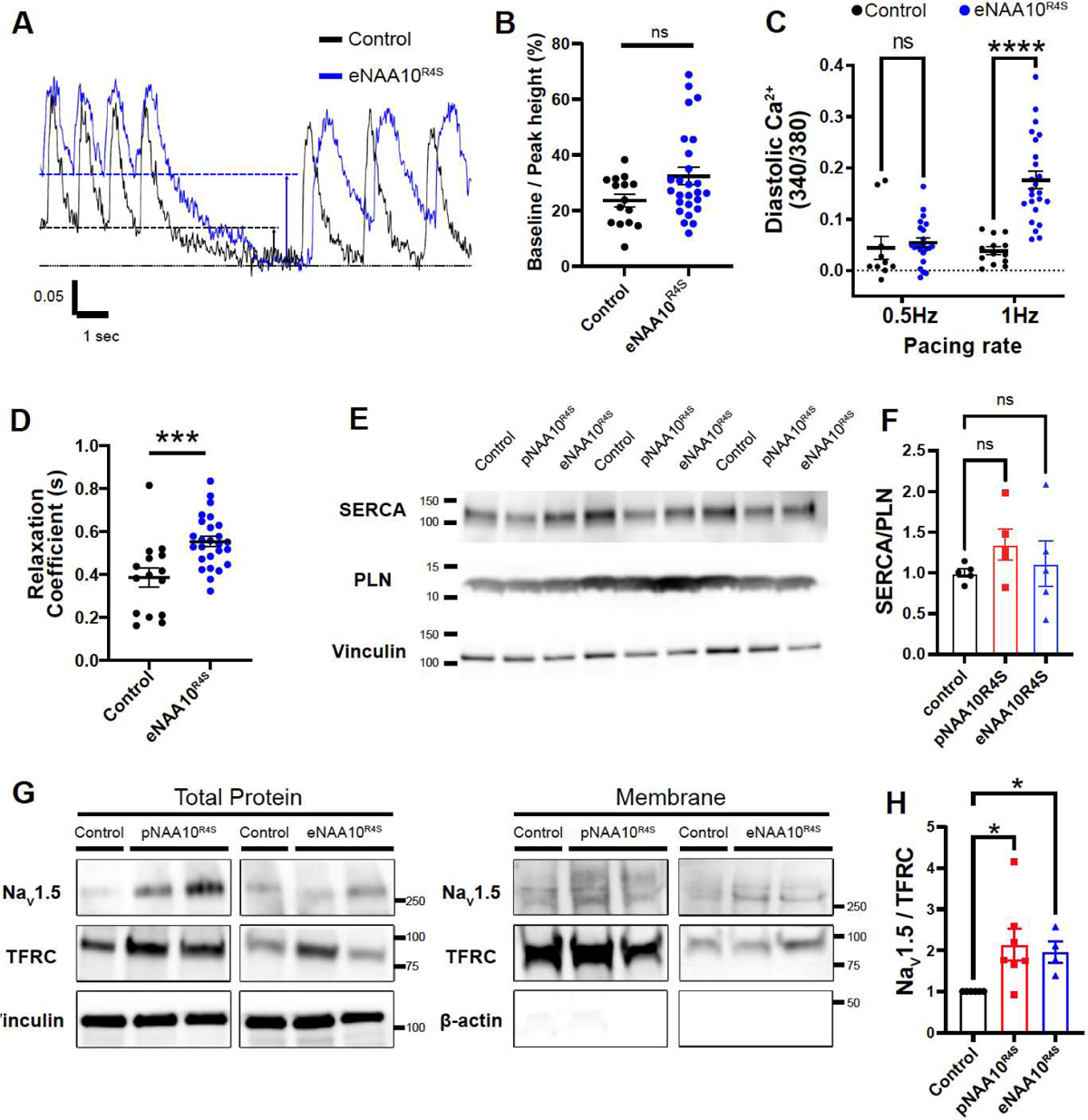
Alterations in Ca^2+^ and Na^+^ handling underlie NAA10^R4S^ cardiomyocyte dysfunction. **A.** Micro-patterned single iPSC-CMs loaded with Fura-2 were electrically paced at 0.5Hz and 1Hz. Diastolic Ca^2+^ levels are indicated by dashed lines. **B.** Quantification of the base to peak height of Ca^2+^ transients paced at 0.5Hz. **C.** End-diastolic Ca^2+^ levels were averaged for five transients and compared to the baseline with electrical pacing at 0.5Hz and 1Hz. **D.** Single exponentials were fit to the relaxation portion of each Ca^2+^ transient to calculate the relaxation coefficient τ. **E.** Whole cell lysates were analyzed by western blot with antibodies specific for SERCA2a (SERCA), phospholamban (PLN) and vinculin. **F.** The ratio of total PLN monomer to SERCA was quantified for each genotype. **G.** Cultures of iPSC-CMs were biotinylated and total and membrane fractions were isolated by streptavidin pull down. Western blot analysis confirms separation of total whole-cell lysates (left panel) and membrane fraction with the expression of the transferrin receptor (TFRC) and lack of b-actin (right panel). **H.** The total membrane fraction of Na_V_1.5 was normalized to TFRC protein levels. Statistics by one-way ANOVA, ****P<0.0001, ***P<0.001, NS > 0.05.

During each Ca^2+^ transient, Ca^2+^ is cleared from the cytosol by re-uptake into the sarcoplasmic reticulum (SR) via the SR Ca-ATPase 2a (SERCA2a) and by Ca^2+^ efflux across the cell membrane via the sodium/calcium exchanger (NCX1)^38^. SERCA2a is inhibited by the binding phospholamban (PLN). To determine if SERCA2a or PLN levels were affected by NAA10 dysfunction, we performed western blotting on whole-cell lysates from control, pNAA10^R4S^ and eNAA10^R4S^ iPSC-CMs. We did not observe significant changes in the level of these proteins or in the ratio of SERCA2a to PLN (Figures 6E and 6F).

Next, we investigated non-SERCA Ca^2+^ efflux, which is influenced by cytoplasmic Na^+^ concentration. Our sodium channel recordings indicated increased activity in mutant NAA10^R4S^ iPSC-CMs (Figures 4A-D). To probe the underlying mechanism, we first determined if the NAA10R4S variant affected SCN5A expression. Quantitative PCR (qPCR) of total RNA from eNAA10^R4S^ iPSC-CMS did not reveal any significant changes RNA transcript levels for SCN5A or other ion channels (Supplemental Figure 4). We next measured the surface expression of Na_V_1.5. After biotinylating surface-accessible proteins of iPSC-CMs, we then pulled down the membrane fraction with streptavidin. Western blot analysis revealed a significant increase in the surface expression of Na_V_1.5 in pNAA10^R4S^ and eNAA10^R4S^ (Figures 6G and 6H), consistent with the increased I_Na_ density by whole-cell patch-clamp (Figures 4A and 4B). These data suggest that NAA10 dysfunction increases Na_V_1.5 activity, inducing Na^+^ overload that increases diastolic Ca^2+^ levels via NCX1.

### Rescue of NAA10 disease phenotypes

In our NAA10-related syndrome iPSC models, decreased NAA10 protein level and catalytic activity correlated with cardiomyocyte dysfunction, suggesting that restoration of NAA10 activity may reverse the disease phenotypes. To test this hypothesis, we generated adenovirus that expresses WT NAA10 along with the self-labeling protein HaloTag (Ad-NAA10, Figure 7A). We transduced cultures of NAA10-mutant iPSC-CMs with either Ad-NAA10 or control adenovirus expressing LacZ (Ad-LacZ) (Figure 7B). Western blot analysis of iPSC-CM lysates 48 hours after transduction demonstrated robust expression over baseline levels (Figure 7C). The endogenous protein migrated at a different size than exogenously expressed NAA10 (Figure 7C, gray arrow).

**Figure 7.**
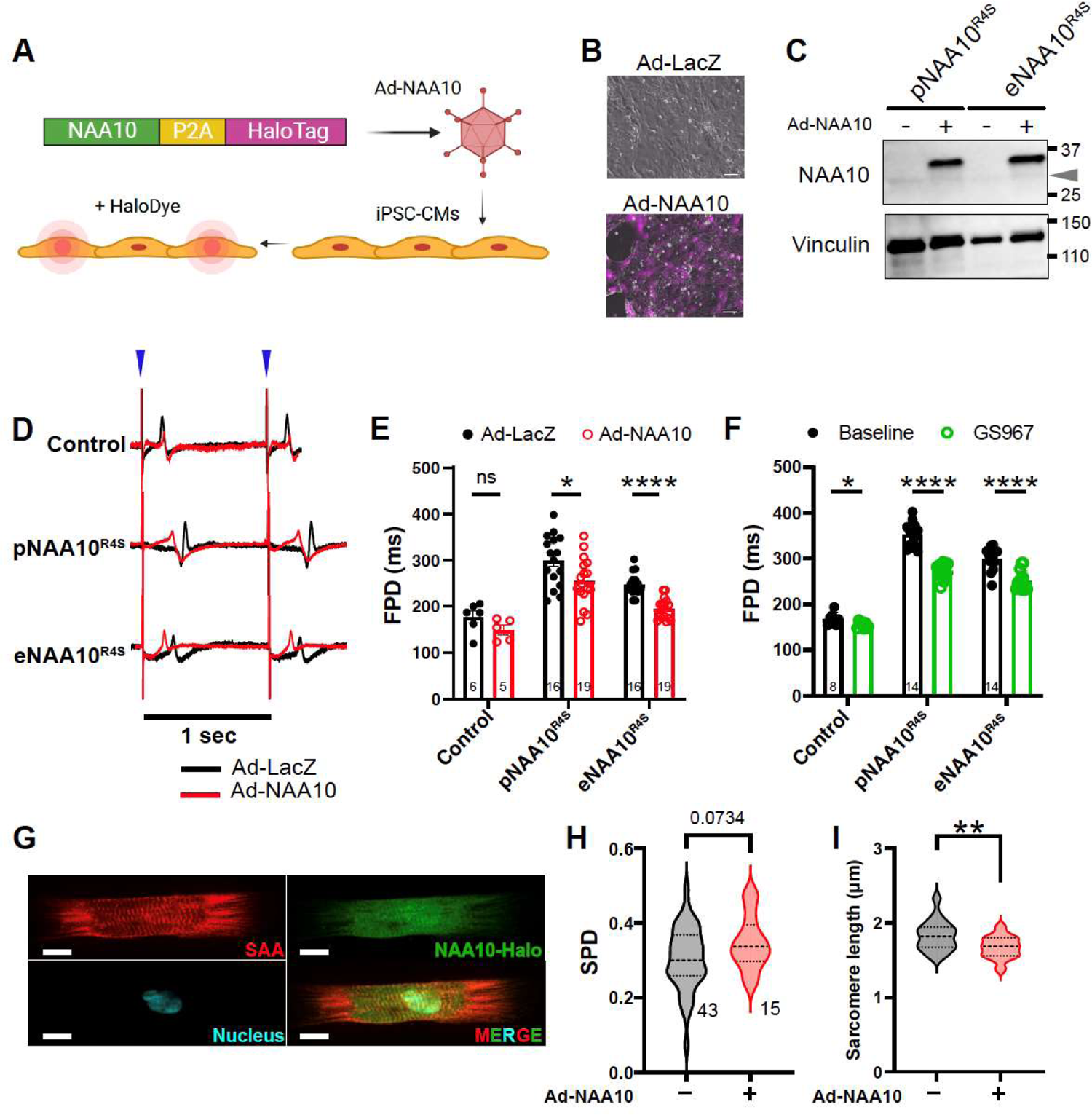
Genetic and pharmacologic rescue of mutant NAA10 iPSC-CMs. **A.** Expression vectors for LacZ or NAA10-P2A-HaloTag were packaged into adenovirus and used to transduce monolayers of iPSC-CMs. **B.** Incubation with HaloTag dye indicates expression of NAA10 (lower panel). Scale bars = 100μm. **C.** Western blot analysis of whole-cell lysates from pNAA10R4S or eNAA10R4S transduced with Ad-LacZ or Ad-NAA10 and probed with anti-NAA10. The faint lower band represents native NAA10 expression (gray triangle). **D.** Representative MEA recordings of monolayers of iPSC-CMs co-transduced with Ad-ChR2-GFP and either Ad-LacZ or Ad-NAA10, plated on MEAs and optically paced at 1Hz (blue triangles). **E.** Quantification of field potential duration (FPD) from D measured as 48 hours after adenovirus transduction. **F.** Monolayers plated on MEAs were first measured at baseline and then treated with the I_NaL_ blocker GS967. The absolute FPD was determined for each genotype and compared as paired treatments. **G.** Single micro-patterned iPSC-CMs transduced with Ad-NAA10 were stained for SAA to quantify changes in sarcomeric organization. Automated analysis of single iPSC-CMs on micro-patterned substrates for **H**. sarcomeric packing density and **I**. Sarcomere length.

Next, we characterized the effect of Ad-NAA10 on NAA10-R4S mutant and wild-type iPSC-CMs. We transduced iPSC-CM monolayers plated on MEAs with either Ad-LacZ or Ad-NAA10 at a multiplicity of infection (MOI) to achieve at least 80% transduction. Cells were also transduced with Ad-ChR2-GFP. Under optical pacing at 1 Hz, Ad-NAA10 treatment did not significantly affect wild-type iPSC-CMs, suggesting that NAA10 overexpression is well tolerated (Figure 7D and 7E). Meanwhile, Ad-NAA10 significantly shortened FPD in NAA10-R4S mutant iPSC-CMs (Figure 7D and 7E).

To determine if NAA10 over-expression could reverse the observed structural defects in NAA10-mutant iPSC-CMs, we plated transduced iPSC-CMs with Ad-NAA10 and plated them on micro-contact printed substrates (Figure 7G). Unbiased computational analysis of confocal micrographs of immunostained iPSC-CMs demonstrated significant shortening of the mean sarcomere length (Figure 7I) and a trend towards increased sarcomeric packing density (Figure 7H) suggesting a partial rescue of structural phenotypes by NAA10 supplement. Collectively these data demonstrate that gene replacement may be a viable therapeutic strategy for NAA10-related syndrome, but additional optimization may be necessary.

To assess the contribution of increased I_NaL_ to the repolarization abnormalities in pNAA10^R4S^ or eNAA10R4S iPSC-CMs, we treated MEA-plated monolayers with the selective late current sodium channel blockers GS967 and ranolazine^39^. GS967 and ranolazine significantly shortened FPDs of NAA10-R4S mutant iPSC-CMs but did not completely normalize them to WT values (Figure 7F; Supplemental Figure 5). In contrast, mexiletine, a class IB anti-arrhythmic which also has effects on potassium channels^40^, did not significantly change the FPD (Supplemental Figure 5).

## DISCUSSION

In this study we created and characterized an iPSC-CM model of NAA10-related syndrome to gain mechanistic insight into the function of NAA10 and how its dysregulation impairs cardiac repolarization and cardiac contraction. By combining optogenetics, cellular electrophysiology, bioengineering, and stem cell biology, our study demonstrates that Nt-acetylation by NAA10 is required for normal regulation of I_NaL_ and I_Ks_, diastolic Ca^2+^, and for normal sarcomere assembly and contraction.

From our clinical cohort, we identified a novel NAA10 variant that segregated with disease phenotype supporting its pathogenicity. A KCNH2 variant was also present in some family members but there is little evidence that this variant is pathogenic, and it was non-segregating. Consistent with other reports of NAA10-related syndrome, we observed variable clinical manifestations in gene-positive patients, with a predominance of SCD in male patients but at least one terminal cardiac event occurring in a carrier female (III-6). One of our patients also developed severe dilated cardiomyopathy, further broadening the cardiac phenotypes associated with NAA10-related syndrome. Like other NAA10 syndrome patients with cardiac involvement, our identified variant is near the extreme N-terminus of NAA10 and within the NAA15 interaction and catalytic domains. Our biochemical data demonstrates that the R4S mutation not only induces NAA10 protein degradation and directly inhibits its Nt-acetylation activity but also destabilizes NAA10’s interaction with NAA15. Our results parallel an early report of NAA10-related syndrome, where the pathogenic variant p.Y43S induced NAA10 protein degradation and impairment of catalytic function^8^. However, the authors did not directly investigate the effects of the Y43S variant on the interaction of NAA10 and NAA15. Interestingly the reported clinical cardiac outcomes were milder without any documented arrhythmia or cardiomyopathy despite QT prolongation. Therefore, our data not only strongly supports that the NAA10 p.R4S is a pathogenic variant but suggests that the disruption of the interaction of NAA10 and NAA15 may be a cause of more severe cardiac involvement.

To investigate the effects of NAA10 dysfunction caused by the R4S mutation, we developed the first iPSC models through somatic-cell reprogramming from affected patients and created an isogenic iPSC line by introducing the same variant into a control line by genome-editing. Our iPSC lines are accurate models of NAA10 dysfunction based on several lines of evidence. First, NAA10 protein levels in mutant iPSC-CMs are reduced by more than 50% consistent with reduced protein stability. Second, NAA10^R4S^ iPSC-CMs have repolarization abnormalities as both monolayers and single cells consistent with QT prolongation. Third, single iPSC-CMs and EHTs derived from NAA10^R4S^ lines had profound defects in sarcomere formation and contractility respectively. These data strongly support that our NAA10-mutant iPSC models accurately recapitulate the clinical phenotype of severe arrhythmogenesis, and cardiomyopathy as seen in our clinical cohort.

We systemically examined each major ionic current underlying the cardiac action potential to identify dysregulation of both I_Na_ and I_Ks_ in mutant NAA10 iPSC-CMs. This unique combination induces significant repolarization abnormalities consistent with a high-risk clinical phenotype (QT > 500msec, recurrent TdP) and similar to patients with complex heterozygosity and missense mutations in both KCNQ1 and SCN5A^41^. In addition to decreased I_Ks_, I_NaL_ was increased as a major mechanism for NAA10-mediated dysregulation of the sodium channel complex. This assertion is supported by recurrent ventricular arrhythmias occurring predominantly during periods of rest and despite high-dose beta-blocker therapy, which are typical clinical features of pathogenic SCN5A variants and LQT3^42^. The contribution of I_NaL_ to abnormal repolarization in NAA10^R4S^ iPSC-CMs is further supported by the shortening of the FPD in response to GS967 and ranolazine, both selective I_NaL_ blockers^39, 43^. In contrast mexiletine, a class 1B sodium channel blocker, failed to shorten the FPD in MEA recordings of eNAA10^R4S^ iPSC-CMs (Supplemental Figure 4). While mexiletine is moderately selective blocker of I_NaL_, it also inhibits the repolarizing K^+^ current I_Kr_ potentially explaining the apparent discrepancy^40^. Mexiletine’s failure to rescue repolarization defects in eNAA10^R4S^ iPSC-CMs, further supports the notion that I_Ks_ current density is reduced by NAA10 dysfunction because it would be the major repolarizing current in the setting of I_Kr_ blockade. Indeed, removal of mexiletine’s off-target effects against I_Kr,_ has been the subject of recent research to improve mexiletine as a selective therapy for LQT3 patients^44^.

In this study, we also propose that NAA10 dysfunction also causes structural and contractile defects in cardiomyocytes. This ascertain is unique, as monogenetic causes of cLQTS are not typically associated with decreased contractility or heart failure^28^. A potential exception to this convention is that select SCN5A variants are associated with DCM and QT prolongation in some patients^45^. Parallel to this association, we detected increased sodium flux through the dysregulation of I_NaL_ in our iPSC models of NAA10 dysfunction. As a potential mechanism for NAA10-mediated contractile dysfunction, we detected elevated diastolic Ca^2+^ levels in NAA10-mutant iPSC-CMs but without any change in the ratio of PLN/SERCA^46, 47^. As an alternative mechanism, increased Na_V_1.5 activity could reverse the function of NCX1, increase diastolic Ca^2+^, and impair relaxation^48^. This multi-factorial mechanism represents a novel model of severe heart failure coupled with repolarization abnormalities that dramatically increases the risk of sudden death.

Our iPSC lines represent the first models of NAA10 dysfunction demonstrating both repolarization and severe contractile abnormalities. A previous report by Tarsha et. al. modeled NAA15 dysfunction in iPSC-CMs but only detected minor contractile impairment^15^. This could represent incomplete inhibition of the NatA complex, or differential targets caused by individual subunit dysregulation. There was no electrophysiologic assessment of NAA15-mutant iPSC-CMs precluding any speculation about similar effects of NAA15 and NAA10 dysfunction on cardiomyocyte action potential. However, given that NAA10-R4S affects both NAA10 function and its association with NAA15, effect severity may correlate with a hierarchy of NatA complex formation > NAA10 function > NAA15 function, but further investigation is necessary.

IPSC disease modeling has dramatically shortened the ark of therapeutic translation. The reproposing of already approved drugs^49^ and development of targeted gene therapies are just two therapeutic development pathways that have benefited from iPSC modeling of cardiac disorders^50^. To determine if NAA10-related syndrome would be amenable to targeted gene therapy, we developed an adenovirus vector to over-express WT NAA10 in our NAA10-mutant iPSC-CMs. With over-expression of human NAA10 protein through adenoviral transduction, there was only partial reversal of the key pathogenic features of NAA10 dysfunction. However, given that Nt-acetylation is a **permanent** co-translational PTM, it is possible that abnormally Nt-acetylated target proteins with longer half-lives continue to circulate in the cardiomyocyte even after normal NatA function is restored. Therefore, the timing and dose of any proposed gene therapy will need to be optimized. Further testing in animal models of NAA10 dysfunction will also be critical for effective translation.

## CONCLUSIONS

Here, we present the first iPSC models of NAA10 dysfunction through the identification of a novel NAA10 variant. We elucidate the underlying mechanisms for the clinical phenotypes of QT prolongation and arrhythmogenesis in NAA10-related syndrome at the cellular and molecular level. We also demonstrate that NAA10 dysfunction negatively impacts cardiomyocyte structure and contractility. Finally, we identify the first cardiac targets for NatA-mediated regulation and establish Nt-aceytlation as an important modulator of cardiomyocyte hemostasis.

## ACKNOWLEDGEMENTS

We would like to thank Fujian Lu for supplying the adenovirus vector, Raul Bortolin for assistance with the bioreactor and Ping-Zhu Zhou for a mammalian expression vector.

