## Supplemental Material for "Dysregulation of N-terminal acetylation causes cardiac arrhythmia and cardiomyopathy"

**EXPANDED METHODS**

**Gene-editing of iPSCs**

For genome-editing, variant specific sgRNAs were designed with CRISPOR^1^, and synthesized using EnGen sgRNA Synthesis Kit (E3322; New England Biolabs, Ipswich, MA, USA). HDR donor template (Integrated DNA Technologies, Coralville, IA, USA) was designed to include synonymous mutations to avoid further digestion by Cas9.

A doxycycline-inducible control iPSC-line (WTC-Cas9) was used for gene-editing as previously described^2^. Briefly, 16 hours before nucleofection, doxycycline was administered at the final concentration of 2 µg/mL. 5 µg of sgRNA and 5 µg of HDR template were transfected into 1 × 10^6^ doxycycline-treated WTC-Cas9 iPSCs using Human Stem Cell Nucleofector Kit (VPH-5012; Lonza Bioscience, Basel, Switzerland) and 4D-Nucleofector (Lonza Bioscience). The next day, the medium was replenished with fresh mTeSR1 without doxycycline, and the single cells were seeded sparsely to a 10cm dish a few days later. Thereafter, colonies were picked up for sequencing. Off-target sites were predicted by CRISPOR^1,3^, and the sequence was analyzed by Sangar sequencing.

**Supplemental Table. PCR primers for amplifying the off-target loci.**

| # | Off-target position | Forward primer | Reverse primer |
| --- | --- | --- | --- |
| 1 | Chr2:129374717 | CAGTGCTGAGTGAAGGCAGA | CCTCTCTGAATTTGGGCCGTC |
| 2 | Chr11:1492482 | ATGAGCGGCAGAAACCAAGA | GCAGCACGCACTTGTGTAAAC |
| 3 | Chr17:75243875 | GCAGGGCGATGTTTTAGCAG | CAGCATCTTCCTTGGAGGGGC |
| 4 | Chr8:59491100 | GTCTTGCAGTTGTGTGGTGC | GAAAGGGAGGCATGCAGCTAT |
| 5 | ChrX:100869663 | TGCCATGGCTTTAGGGAAGG | CCGGCACAGAGAAATCCCTTC |

**Differentiation and purification of iPSC-CMs**

IPSCs were differentiated into CMs as previously described^4^. Briefly, 2 days after iPSCs were seeded onto a 12-well plate, the medium was replaced with RPMI / B27 minus insulin (Thermo Fisher Scientific, Waltham, MA, USA) containing 6 µM CHIR99021 (Stemcell technologies, Vancouver, Canada) (Day 0). On Day 2, the medium was replaced with RPMI/B27 minus insulin containing 5 µM IWP2 (Tocris Bioscience, Bristol, UK). On Day 4, the medium was replaced with RPMI/B27 minus insulin. On Day 6 and every 3 to 4 days thereafter, RPMI/B27 medium was replenished. CMs were purified in glucose-depleted lactate-supplemented medium as previously described^5^. Differentiated CMs on Days 50 through 90 were used for experiments.

WTC-11 (Ctrl) is a wild-type human male iPSC line (Coriell Institute: # GM25256) that harbors a doxycycline (Dox)-inducible CRISPR/Cas9, which was created as described previously^6^ and cultured in E8 medium (Life Technologies, # A1517001). IPSCs were differentiated in the DASGIP® Parallel Bioreactor Systems (Eppendorf) as described previously^7^ with several modifications that will be part of an independent manuscript. In short, hiPSCs were cultured in T80 cell culture flasks (Life Technologies, # 178905) pretreated with 1:100 (v/v) diluted Geltrex (Life Technologies, # A1413302) at an initial seeding density of 15,000 cells/cm^2^. Cells were maintained in E8 medium (Life Technologies, # A1517001) with daily medium change until 90% cell confluency was reached. For dissociation, T80 flasks were washed once with PBS, incubated with 5 mL Versene (Life Technologies, # 1540066) for 15 to 20 min at 37°C and dissociation was stopped by adding 5 mL E8 medium. 50 million single iPSCs were resuspended in 50 mL E8 medium supplemented with 10 μM of ROCK inhibitor. The bioreactor vessel was taken from the bioreactor system and placed under the laminar flow and 50 mL E8 (+ 10 μM Y-27632) and 50 mL of the iPSC solution resulting in a final volume of 100 mL per vessel. Cells were agitated at a speed of 60 rpm, passed with 21% O_2_ and 5% CO_2_ by 10 sL/h overlay gassing and maintained at 37°C. The next day diameter of spontaneously formed embryoid bodies (EBs) was measured to estimate time of differentiation start. If critical diameter (100-300 µm) was reached, cardiac differentiation was induced by a complete change of the medium to RPMI 1640 with B27 (w/o Insulin; basic medium) supplemented with 7 μM CHIR99021 (day 0). After 24 h (Day 1), the complete medium was changed to basic medium, and cells were incubated for an additional 24 h. On day 2, the complete medium was changed again to basic medium containing 5 µM IWR-1-endo for 48 h. On day 4, the complete medium was changed to basic medium, and cells were incubated for an additional 72 h. From day 7 on cells were cultured in basic medium supplemented with 1:1000 (v/v) insulin (Sigma-Aldrich, # I9278) followed by 50% medium refreshments at day 9, 11 and 13. Finally, cells were dissociated on day 15 for 3-4 h depending on EB size and density using Collagenase II (Worthington, # LS004176)^8^, and frozen using a controlled rate freezer (Grant, CRF-1).

**Electrophysiology**

**Whole-cell patch clamp recordings**

Cultured iPSC-CMs were dissociated with Accutase and plated sparsely onto Geltrex-coated 11mm coverslips. Single iPSC-CMs were analyzed 3 to 6 days after dissociating. Single iPSC-CMs were recorded under different conditions to acquire each parameter (Supplemental Table 1)^910^. Perforated patch recordings were performed for action potential (AP) analysis and the L-type Ca^2+^ current (I_CaL_). The ruptured patch technique was used for I_Na_, I_Ks_, and I_Kr_ recordings. For AP analysis, iPSC-CMs exhibiting a APD90/APD50 ratio of less than 1.4 were defined as ventricular type^11^. I_NaL_, I_Ks_, and I_Kr_ were defined as currents specifically sensitive to 30 µM TTX, 1 µM HMR1556, and 1 µM E4031 respectively. The current traces were subtracted before and after the drug administration to elicit those specific currents. Pipettes were pulled from thick-walled borosilicate glass capillaries (1B150F-4; World Precision Instruments, FL, USA) for AP and I_CaL_, and from thin-walled capillaries (TW150-4; World Precision Instruments) for I_Na_, I_Ks_, and I_Kr_. dPatch^®^, and SutterPatch^®^ (Sutter Instrument, CA, USA) were used for data acquisition. All the data were acquired from at least three independent experiments using different biological replicates.

**Multi-electrode array with optogenetics**

Single iPSC-CMs were isolated by incubating collagenase-B (Roche, Roswell, GA, USA, 1mg/mL) for 15 minutes and thereafter 0.25% Trypsin or Accutmax (Innova Cell Technologies, San Diego, CA, USA) for 5 minutes for the dissociation as previously described^11,12^. Cell suspensions (3 × 10^4^ cells in 5 µL) were placed onto fibronectin-coated multi-electrode array (MEA) plates (CytoView MEA; Axion BioSystems, Atlanta, GA, USA). After 3 days, the CMs were infected with the crude adenovirus expressing Channelrhodopsin-2 fused to green fluorescent protein (GFP) (Ad-ChR2-GFP) to enable optical stimulation. Four or more days after the infection, field potentials (FP) were recorded using Maestro Edge (Axion BioSystems). FP signals were digitally sampled at 12.5 kHz and the system bandwidth is 0.01 Hz – 5 kHz. iPSC-CMs were stimulated by using a multi-well light stimulation system (Lumos 24; Axion BioSystems). Specifically, CMs were stimulated at the rate of interest (1Hz, 2Hz, or 3Hz) for 40 beats and the final 30 beats were averaged. FP duration (FPD) was defined as the interval between a spike and a subsequent positive deviation. This parameter was automatically measured with Cardiac Software Module on the system. All the data were acquired from at least three independent biological replicates.

**Immunofluorescence**

Samples were washed with cold Ca^2+^-free phosphate buffered saline (PBS) for 5 minutes before fixed with 4% paraformaldehyde for 10 minutes at 4°C, and permeabilized with 0.1% Triton X in PBS for 10 minutes at room temperature. Blocking was performed with 3% bovine serum albumin in PBS. Primary antibodies and secondary antibodies were sequentially incubated for 2 hours at room temperature, followed by serial washes with PBS and DAPI in the mounting media. The details of antibodies are listed in supplemental table. The coverslips with stained samples were mounted with Prolong Diamond Antifade Mount (Invitrogen, 2273639), and after 24 hours of incubation in dark conditions at room temperature, imaging was performed. Confocal microscopy (Olympus FV3000R) with a 60x oil immersion objective was used.

| Primary antibodies | | | |
| --- | --- | --- | --- |
| Antigen | Manufacturer | Catalog number | Concentration |
| NAA10 | Atlas antibodies | HPA030711 | 1:1000 (WB) |
| ARD1A | Proteintech | I4803-I-AP | 1:3000 (WB) |
| NAA15 | Atlas antibodies | HPA023589 | 1:1000 (WB) |
| Vinculin | Santa Cruz | sc-73614 | 1:200 (WB) |
| β-Actin | Cell Signaling Technology | 3700 | 1:1000 (WB) |
| SCN5A | Alomone labs | ASC-005 | 1:500 (WB) |
| DDDDK tag (FLAG tag) | Abcam | ab1257 | 1:1000 (WB) |
| Transferrin receptor | Invitrogen | 13-6800 | 1:1000 (WB) |
| SERCA | Santa Cruz | sc-376235 | 1:100 (WB) |
| PLN | Cell Signaling Technology | 14562 | 1:1000 (WB) |
| α-Actinin (Sarcomeric)  (FITC-conjugated) | Miltenyi Biotec | 130-119-766 | 1:50 (IF) |
| Phalloidin  (Alexa647-conjugated) | Invitrogen | A22287 | 1:400 (IF) |
| WGA  (Alexa647-conjugated) | Invitrogen | 2126807 | 1:1000 (IF) |
| Secondary antibodies | | | |
| HRP goat anti-mouse IgG | Bio Rad | 51782504 | 1:10000 (WB) |
| HRP donkey anti-rabbit IgG | Bio Rad | 644005 | 1:10000 (WB) |
| IRDye800 CW donkey anti-mouse | LI-COR Biosciences | 926-32212 | 1:10000 (WB) |
| IRDye680 RD donkey anti-goat | LI-COR Biosciences | 926-68074 | 1:10000 (WB) |
| Alexa647 donkey anti-rabbit | Invitrogen | A32795 | 1:1000 (IF) |
| Hoechst 33258 | Molucular Probes | H-3569 | 1:500 (IF) |

**Micro-contact patterning and sarcomere analysis**

Using Computer-Aided Design (CAD), the single-cell pattern was first designed as a series of rectangles with 7:1 aspect ratio (105 μm by 15 μm) surrounded by a 220 μm thick boundary. The design was then converted to photolithography masks (CAD/Art Services Inc.). At the Center for Nanoscale Systems (Harvard University), silicon wafers (Wafer World) with a diameter of 3 inches are cleaned with a nitrogen gun and then spincoated with photoresist SU8-3005 (MicroChem Corp.), followed by cycles of 1 min and 2 minutes of baking at 65 and 95 degrees Celsius, respectively. Polymerization via UV-light exposure through the photomasks with the desired single-cell patterns is then executed for 20 seconds. The cycles of baking are repeated, and the post-UV wafers are developed in propylene glycol methyl ether acetate (PGMEA, Sigma) for no more than 1 minute under vigorous agitation. After development, the wafer is desiccated overnight with a small amount of silane (United Chemical) to prevent Polydimethylsiloxane (PDMS) from binding to it permanently. Once the wafers are prepared, Polydimethylsiloxane (PDMS, Sylgard 184; Dow Corning) prepared at a ratio of 10:1 (base: curing agent) is placed on the wafer, covering its entire surface, and baked for 24 hours at 65 degrees Celsius. The next day, stamps are cut without damaging the wafer and sonicated in 70% ethanol for use in patterning.

Glass coverslips (12 mm, VWR, 48366-252) were spin-coated at custom recipes with a 1:1 ratio of Polydimethylsiloxane (PDMS) elastomer (Sylgard 184, The Dow Chemical Company, Midland, MI, USA) and dielectric gel (Sylgard 527, The Dow Chemical Company). The latter, PDMS 527, is prepared via combining Part A and Part B of the kit in a 1:1 ratio. Coverslips were coated for 48 hours in a 65°C oven. Thereafter, stamps containing rectangular-shaped islands with 7:1 aspect ratio^13^ were first coated for 1 hour with Fibronectin (Sigma Aldrich F0895) (50 µg/ml) diluted in Geltrex (1:200, Life Technologies, A1413302). Fibronectin aliquots of 1 mg/ml were prepared in phosphate-buffered saline (PBS) and stored at -20°C. In the meantime, PDMS-coated coverslips are exposed to UV Ozone (Jelight) for 8 minutes and the patterning process was performed by placing the dried stamps onto the coverslips. The coverslips were immersed in 1% Pluronic F-127 (Sigma-Aldrich, P2443) for less than 10 minutes to block the portions of the coverslips not coated with fibronectin, followed by washing 3 times with room-temperature PBS.

iPSC-CMs were seeded onto the micropatterned coverslips three days before immunofluorescence imaging. Cells were stained with FITC-conjugated α-sarcomeric actinin, and Alexa-647 conjugated Phalloidin, followed by staining with Hoechst 33342.

To assess sarcomere alignment in micropatterned iPSC-CMs, we used an unbiased algorithm developed by the Disease Biophysics Group incorporating the ImageJ Plugin (Orientation J) and a custom-made MATLAB (Mathworks)script for structural analysis of single cells^14^. Briefly, a Sarcomere Packing Density (SPD) reflects the degree of spatial organization of the sarcomeres quantifying the immunosignal localized in a regular lattice and the periodicity of the positive structures respectively of their orientations. This means that the poorly formed sarcomeres that are not periodically spaced demonstrates reduced SPD value, ranging from 1 to 0.

**EHT generation**

Engineered heart tissues (EHTs) were made as previously described^8^. Briefly, 3D differentiated iPSC-CMs were thawed gently, and 0.8 × 10^6^ cells were resuspended in 100 µL of B27/RPMI medium containing 10 µM Y-27632 (TOCRIS, 1254), 10% Matrigel (Corning, 354234), and 5 mg/mL fibrinogen (Sigma, F8630) for a single EHT. Immediately after adding 300 mU thrombin, the cell suspension was transferred into a 2% agarose mold where racks containing silicone pillars were embedded in a 24-well plate and incubated at 37°C for 1.5 hours. Thereafter, the silicon racks were transferred to a new plate with 1.5mL RPMI/B27 supplemented with aprotinin (Sigma, A1153).

Engineered heart tissues (EHTs) were generated as described previously^5,42^ with some minor modifications. Briefly, 0.8x10^6^ hiPSC-CMs were used to generate each EHT. Cells were transduced with adenovirus ChR2-YFP on the day of casting or after 7 days in vitro. We modified the standard EHT culture medium (EHT-medium in the referenced literature^5^) by replacing DMEM with RPMI 1640 plus B27 minus insulin, removing 10% heat-inactivated horse serum, and reducing aprotinin concentration to 5 µg/ml. This resulted in EHTs initiating contraction as early as day 1-3 after EHT assembly, versus day 7-10 reported in the literature^5^. EHT contraction was recorded as described below from day 7 on and functional analysis was performed from day 27 to day 33. For Immunohistochemical analysis EHTs were cryosectioned and stained with primary antibodies shown in Table S4. Sections were imaged using an Olympus FV-3000 confocal microscope and analyzed with Fiji (ImageJ).

***Functional assessment of EHTs***

EHTs in a 24 well plate were placed in a stage top incubator and maintained at 37C, 5% CO2. EHTs were optically paced at different frequencies using blue LEDs positioned above the place and recorded from below at 30 frames per second through a 561 nm long-pass filter (Semrock BLP02-561R-32) using an 8mm f/1.4 lens (ThorLabs MVL8M1) mounted on a Basler acA1920 camera. EHT post movement was tracked post-hoc using the multi-template matching FIJI plugin^43^. Twitch force measurements were subsequently measured by applying post deflection to the beam bending theory for a known Young’s modulus of the posts, as described in detail elsewhere^44^.

**Quantitative PCR**

Cells were washed once with ice-cold PBS and lysed in TRIzol. Total RNA was extracted by centrifugation and RNA samples were isolated. Coding DNA (cDNA) was made using a reverse transcriptase kit (Superscript III, Invitrogen). We quantified total cDNA for each sample and normalized the concentration. Quantitative PCR was performed on a 96 well thermocycler (BioRad) at an annealing temperature of 55C with validated gene-specific primers. Ct values were compared to a house-keeping gene (GAPDH) and the fold-change was calculated and compare to control samples for each gene transcript.

**Western blot**

Cells were lysed with mTOR lysis buffer (120mM NaCl, 40mM HEPES, 40mM NaF, 1mM EDTA, 10mM β-Glycerophosphate disodium, 0.3% CHAPS, pH 7.5 with NaOH) containing 1% TritonX and Halt protease and phosphatase inhibitor (Life Technologies 78442). The concentration of the protein was measured with BCA protein Assay kit (Thermo scientific 23225) and 10 µg of protein in each lane was analyzed by SDS-PAGE and immunoblotting. Blots were incubated with primary antibodies and secondary antibodies sequentially for 2 hours at room temperature or overnight at 4℃. Protein signals were detected using an enhanced chemiluminescent substrate (BioRad), and images were captured using Azure 300 (Azure Biosystems, Dublin, CA, USA) and analyzed with ImageJ. The antibodies are listed in supplemental table.

**Expression plasmids**

NAA10^WT^-3×FLAG cloned into pUC-GW-Kan vector was synthesized by a manufacturer (GENEWIZ, South Plainfield, NJ, USA). Then, NAA10 ^WT^ -3xFLAG was cloned into pcDNA3.1(+) vector, and NAA10^R4S^-3xFLAG/pcDNA3.1(+) was generated by site-directed mutagenesis with In-Fusion cloning (Takara Bio, Kusatsu, Japan).

NAA15 expression plasmid (HG19640-UT, Sino Biological) was also cloned into pcDNA3.1(+) vector and myc tag was added using In-Fusion cloning.

**Cycloheximide chase experiment**

2.5 µg of plasmids were transfected into HEK293T cells on a 6-well plate with Lipofectamine 3000. 48 hours after the transfection, the medium of each well was replaced with 2 ml culture medium containing 50 µg/mL Cycloheximide (Sigma, 01810). Cells were harvested at 0, 2, 4, and 6 hours after the Cycloheximide administration. The cells harvested at 0 hour were not treated with Cycloheximide. Harvested cells were centrifuged at 4°C for 15 seconds with 14000g, and the cell pellets were washed with ice-cold PBS, centrifuged again, and stored at -80°C after the supernatant was discarded. After all the samples were harvested, western blot was performed, and the membrane was stained with anti-DDDDK tag antibody (Abcam, ab1257) and anti-Vinculin antibody (Santa Cruz). For secondary antibodies, IRDye680 RD donkey anti-goat (LI-COR Biosciences, Lincoln, NE, USA) and IRDye800 CW donkey anti-mouse (LI-COR Biosciences) were used. Imaging was performed with LI-COR Odyssey Infrared Imaging System (LI-COR Biosciences).

**NAA10-NAA15 binding assay**

The NAA10- and NAA15-expression plasmid was transfected to HEK293T cells with Lipofectamine3000. 48 hours later, the cells were harvested and lysed with mTOR lysis buffer. The lysate was incubated with anti-FLAG magnetic beads overnight at 4°C. The protein was eluted with FLAG peptide and analyzed with SDS-PAGE. Immunoblotting was performed as described above. The protein was quantified using ImageJ.

**Protein synthesis**

50 µg of the NAA10 expression plasmid was transfected into HEK293T cells on a 15 cm dish using polyethylenimine (PEI) (Sigma 408727). 4 dishes were prepared for each plasmid, and after lysing the cells the cell lysate was filtered with a 0.22µm filter, and pre-cleared with mouse IgG agarose. Thereafter, FLAG-tagged NAA10 was pulled down with FLAG tag using anti-FLAG magnetic beads (Sigma M8823). The protein was eluted with FLAG peptide and concentrated using Amicon Ultra-4 Centrifugal Filter Unit (Millipore, UFC8010).

**ThioGlo4 assay**

ThioGlo4 assay was modified from a previous protocol^15^. The custom peptide as a substrate for NAA10 (EEEIA24: EEEIAALRWGRPVGRRRRPVRVYP) was synthesized by Biomatik (Kitchener, Ontario, Canada). Briefly, the mixture of 15 µM ThioGlo4 (MilliporeSigma, 59550410MG), 150 mM NaCl, 25 mM HEPES (pH 7.5), and 0.001 % TritonX was reacted with 0.5, 1, 2, 5, and 10 µM of CoA (Sigma, C4780), respectively, for standard curve. Besides, 15 µM ThioGlo4, 50 µM AcCoA (Sigma, A2181), 50 µM Substrate (EEEEIA24), 150 mM NaCl, 25 mM HEPES (pH 7.5), and 0.001% Triton X was reacted with 25, 50, 100, 200 nM of purified NAA10, respectively. All the experiments were performed at 25 ℃ in duplicate, and the fluorescence intensity was measured with an excitation wavelength of 400 nm, and an emission wavelength of 465 nm using a FlexStation^®^3 multi-mode microplate reader. The fluorescence was continuously recorded for 30 minutes with an interval of 30 seconds. The baseline was subtracted, and Vmax was calculated using SoftMax Pro Software.

**Adenovirus generation**

NAA10-P2A-HaloTag was cloned based on pH6HTC His_6_HaloTag^®^ T7 Vector (Promega, Madison, WI, USA). The NAA10-P2A-HaloTag sequence was inserted into pENTR/D-TOPO vector (Invitrogen) and thereafter into pAd/CMV/V5/DEST vector (Invitrogen) following the manufacturer’s protocol. The vector was digested with Pac I, and the Pac I-digested vector was transfected to HEK293A cells with Lipofectamine 3000 on a 6-well plate. Thereafter, the virus was amplified by infecting HEK293A cells on a 10-cm dish. The crude stock was used in experiments and is designated at Ad-NAA10.

**Ca^2+^ imaging**

Single iPSC-CMs were seeded on PDMS-coated micro-patterned coverslips. After 3 days, the coverslips were incubated with 5 µM Fura-2 (ThermoFisher, F14185) at 37℃ with 5% CO2 for 20 minutes, and, after washing, placed in a C-Stim CMC microscope chamber (IonOptix) and a temperature of 36-37°C was maintained by using a mTCII Temperature Controller (IonOptix) to circulate extracellular by a closed-loop controller. Extracellular buffer containing (in mM) NaCl 140, KCl 5.4, MgCl2 1.2, CaCl2 1.8, HEPES 10, Glucose 10, and sodium pyruvate 2, with pH of 7.4, was used in this experiment. The samples were imaged using IonOptix Calcium Imaging system installed on an Olympus IX71. During the imaging, the cells were stimulated by MyoPacer (IonOptix). The background was subtracted, and the data were analyzed using IonWizard software (IonOptix).

**Cell-surface Biotinylation**

After washing with PBS (+) (PBS containing 0.2mM CaCl_2_ and 1.5mM MgCl_2_), cells were incubated with 0.5 mg/mL Sulfo-NHS-SS-Biotin (Thermo) in PBS(+) for 1 hour on ice with occasional shaking. Thereafter, the biotinylating reaction was quenched by washing 3 times with PBS(+) containing 100 mM glycine. Cells were lysed with mTOR buffer, and part of the lysate was saved as total protein. The rest of the lysate was incubated with Pierce Streptavidin Magnetic Beads (Thermo) at 4 °C overnight on a rotator. The magnetic beads were collected on a magnetic stand, and the protein sample was eluted by incubating with SDS-PAGE reducing sample buffer at 96-100 °C for 5 minutes.

**Bioinformatics**

A multiple sequence alignment of NAA10 sequence was performed with Clustal Omega^16^, and the annotation of conservation was conducted with Jalview^17^. DynaMut2 was used to predict the impact of the mutation on protein dynamics and stability^18^. Information of protein structure was obtained from the PDB database, and the human NatA amino-terminal acetyltransferase complex (PDB code: 6C9M)^19^, was applied to the analysis.

**Statistics**

Prism 9 (GraphPad, San Diego, CA, USA) was used for statistical analysis. Normal distribution was tested with Shapiro-Wilk test. For normally distributed samples, data were presented as mean ± SEM. An unpaired Student’s t-test or one-way analysis of variance (ANOVA), followed by Dunnett’s comparison test, was used for two- or more than two-group comparisons. For repetitive measurements, a two-way repeated measures ANOVA, followed by Dunnett’s comparison test, was performed. For samples without normal distribution, data were presented with violin plots. Mann-Whitney test or Kruskal-Wallis test, followed by Dunn’s multiple comparisons test, was conducted for two- or more than two-group comparisons. Results were considered statistically significant at p < 0.05.

**SUPPLEMENTAL TABLES**

Supplemental Table 1. Single-cell electrophysiology recording conditions.

|  | AP | I_Na_ | I_CaL_ | I_Ks_ | I_Kr_ |
| --- | --- | --- | --- | --- | --- |
| Configuration | Current clamp | Voltage clamp | Voltage clamp | Voltage clamp | Voltage clamp |
| Sampling rate (kHz) | 50 | 100 | 50 | 10 | 10 |
| Filter (kHz) | 10 | 20 | 10 | 1 | 1 |
| Temperature (°C) | 35 ± 1 | 20 ± 1 | 34 ± 1 | 34 ± 1 | 34 ± 1 |
| Perforated /Ruptured | Perforated | Ruptured | Perforated | Ruptured | Ruptured |

| Patch Clamp Extracellular Solution (mM) | | | | | | |
| --- | --- | --- | --- | --- | --- | --- |
|  | AP | I_Na_ | I_Na_-Late | I_CaL_ | I_Ks_ | I_Kr_ |
| NaCl | 150 | 70 | 150 | 140 | 150 | 150 |
| NMDG |  | 70 |  | - |  |  |
| KCl | 5.4 | - | 5.4 | - | 5.4 | 5.4 |
| CsCl_2_ |  | 5.4 |  | 10 |  |  |
| CaCl_2_ | 1.8 | 1.8 | 1.8 | 1.8 | 1.8 | 1.8 |
| MgCl_2_ | 1 | 1.2 | 1 | 1 | 1 | 1 |
| Glucose | 15 | 10 | 15 | 10 | 15 | 15 |
| HEPES | 15 | 10 | 15 | 10 | 15 | 15 |
| Na-Pyruvate | 1 | 2 | 1 | - | 1 | 1 |
| TTX | - | - | - | - | - | - |
| Nifedipine | - | 0.01 | 0.01 | - | 0.002 | 0.002 |
| E4031 | - | - | - | - | 0.001 | - |
| HMR1556 | - | - | - | - | - | - |
| pH 7.4 | NaOH | HCl | NaOH | NaOH | NaOH | NaOH |

| Patch Clamp Intracellular Solution (mM) | | | | | |
| --- | --- | --- | --- | --- | --- |
|  | AP | I_Na_ / I_Na_-Late | I_CaL_ | I_Ks_ | I_Kr_ |
| KCl | 150 | - | - | 20 | 150 |
| L-Aspartic Acid | - | 120 | - | - | - |
| CsCl_2_ | - | 20 | 120 | - | - |
| CsOH | - | 120 | - | - | - |
| NaCl | 5 | - | - | - | 5 |
| CaCl_2_ | 2 | - | - | - | 2 |
| EGTA | 5 | 10 | 10 | 10 | 5 |
| HEPES | 10 | 10 | 5 | 5 | 10 |
| MgATP | 5 | - | 5 | 5 | 5 |
| Na2-ATP | - | 4 | - | - | - |
| K-Aspartate | - | - | - | 125 | - |
| MgCl_2_ | - | 2 | - | 1 | - |
| Na_2_-Phosphocreatine | - | - | - | 2 | - |
| Na_2_-GTP | - | - | - | 2 | - |
| Escin | - | - | - | - | - |
| pH | 7.2 (NaOH) | 7.3 (CsOH) | 7.2 (CsOH) | 7.2 (NaOH) | 7.2 (NaOH) |

| Supplemental Table 2 Gating Parameters of Nav1.5 and Cav1.2 in Control- and NAA10^R4S^-iPSC-CMs | | | | |
| --- | --- | --- | --- | --- |
| I_Na_ parameters | Control | pNAA10^R4S^ | eNAA10^R4S^ | p Value |
| Peak I_Na_ density | -103.1 ± 9.8  (n = 14) | -271.6 ± 19.8^**^  (n = 8) | -249.5 ± 30.8^**^  (n = 10) | 0.0017 |
| Steady-state activation  V_1/2_  *k* | -21.6 ± 0.7  6.5 ± 0.2 | -26.8 ± 1.2^*^  6.5 ± 0.2 | -25.8 ± 0.8^*^  6.9 ± 0.6 | 0.0275  0.6465 |
| Steady-state inactivation  V_1/2_  *k* | -69.0 ± 0.9  6.7 ± 0.1 | -77.4 ± 0.8^**^  7.0 ± 0.2 | -75.6 ± 0.8^**^  6.6 ± 0.1 | 0.0010  0.4684 |
| I_Ca-L_ parameters | Control | pNAA10^R4S^ | eNAA10^R4S^ | p Value |
| Peak I_Ca-L_ density | -7.0 ± 1.2  (n = 3) | -6.4 ± 0.5  (n = 8) | -9.8 ± 1.1  (n = 6) | 0.1492 |
| Steady-state activation  V_1/2_  *k* | -6.8 ± 2.4  8.7 ± 0.8 | -10.2 ± 0.8  6.9 ± 0.4 | -10.9 ± 1.6  6.7 ± 0.4 | 0.3246  0.0918 |
| Steady-state inactivation  V_1/2_  *k* | -40.1 ± 1.5  7.9 ± 1.8 | -41.5 ± 1.6  6.6 ± 0.1 | -46.9 ± 0.9  6.5 ± 0.6 | 0.0317  0.8453 |

p Values were calculated using one-way ANOVA or Kruskal-Wallis test.

* p < 0.05, **p < 0.01; Dunnet’s multiple comparisons test for pNAA10^R4S^ or eNAA10^R4S^ versus Control.

| Supplemental Table 3. Parameters of Ca^2+^ transient in Control- and NAA10^R4S^-iPSC-CMs | | | |
| --- | --- | --- | --- |
| Ca^2+^ release | Control (n = 15) | eNAA10^R4S^ (n = 26) | p Value |
| Baseline / Peak height | 23.6 ± 2.27 | 32.4 ± 3.11 | 0.1652 |
| Departure velocity | 3.3 ± 0.60 | 4.2 ± 0.62 | 0.3799 |
| Time to peak | 0.22 ± 0.023 | 0.24 ± 0.016 | 0.4106 |
| Peak height | 0.13 ± 0.011 | 0.21 ± 0.021 | 0.0286* |
| Peak | 0.73 ± 0.026 | 0.85 ± 0.034 | 0.0156* |
| Time to peak 50 | 0.049 ± 0.0093 | 0.049 ± 0.0055 | 0.7330 |
| Time to peak 75 | 0.096 ± 0.012 | 0.091 ± 0.0089 | 0.7349 |
| Time to peak 90 | 0.15 ± 0.018 | 0.15 ± 0.012 | 0.9577 |
| Time to maximal departure velocity | 0.038 ± 0.010 | 0.016 ± 0.0038 | 0.0617 |
| Area of the departure phase | 0.021 ± 0.0027 | 0.038 ± 0.0045 | 0.0099** |
| Ca^2+^ uptake | Control | eNAA10^R4S^ | p Value |
| Return velocity | -0.46 ± 0.054 | -0.54 ± 0.062 | 0.6542 |
| Time to baseline 50 | 0.45 ± 0.031 | 0.54 ± 0.015 | 0.0072** |
| Time to baseline 75 | 0.59 ± 0.038 | 0.70 ± 0.011 | 0.0257* |
| Time to baseline 90 | 0.74 ± 0.042 | 0.84 ± 0.0088 | 0.0701 |
| Time to maximal return velocity | 0.35 ± 0.029 | 0.43 ± 0.022 | 0.0352* |
| Area of the return phase | 0.035 ± 0.0046 | 0.064 ± 0.0058 | < 0.0001**** |

p Values were calculated using unpaired t test or Mann-Whitney test. * p < 0.05, **p < 0.01, ***p < 0.001, ****p < 0.0001

**SUPPLEMENTAL FIGURES AND LEGENDS**

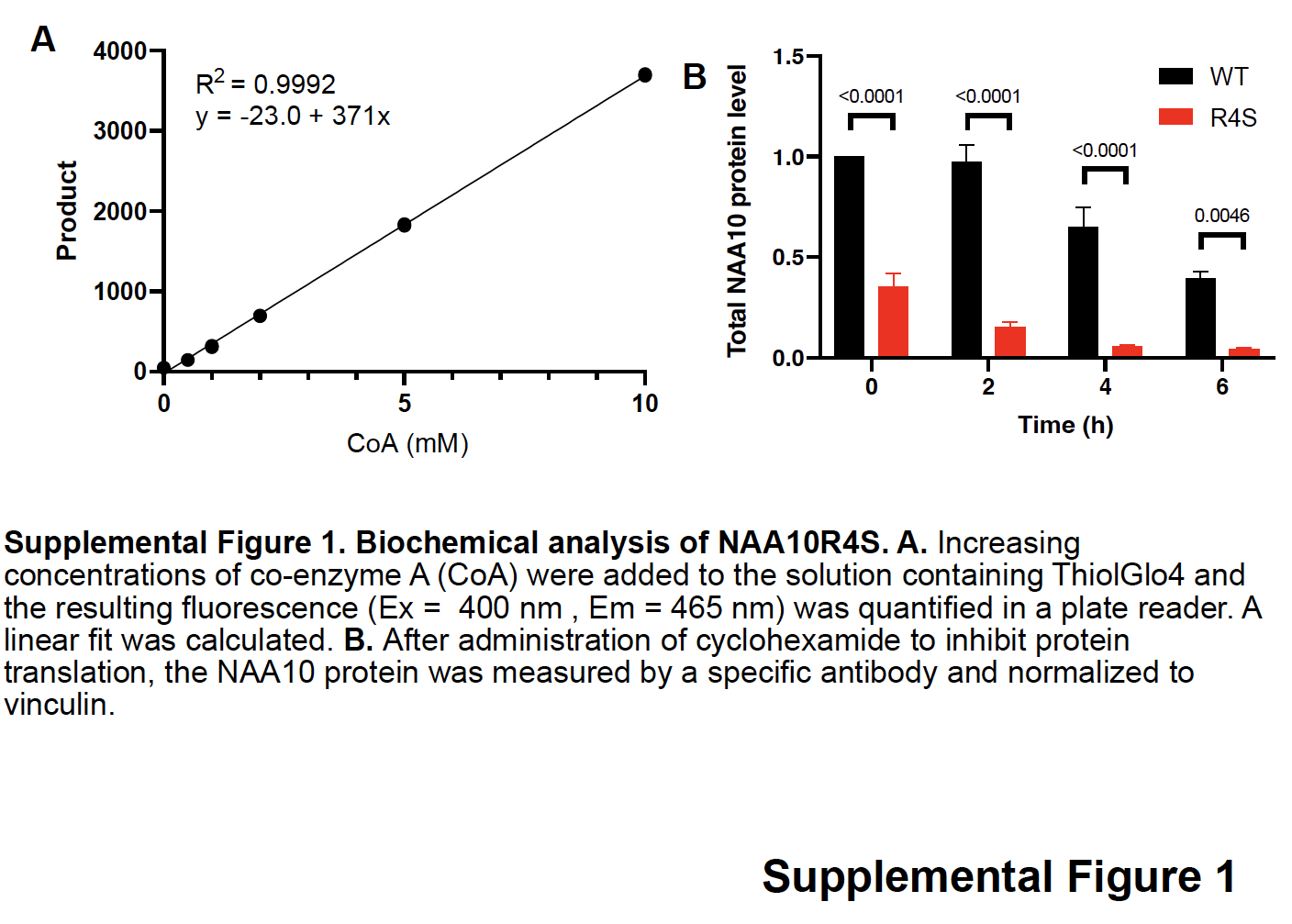

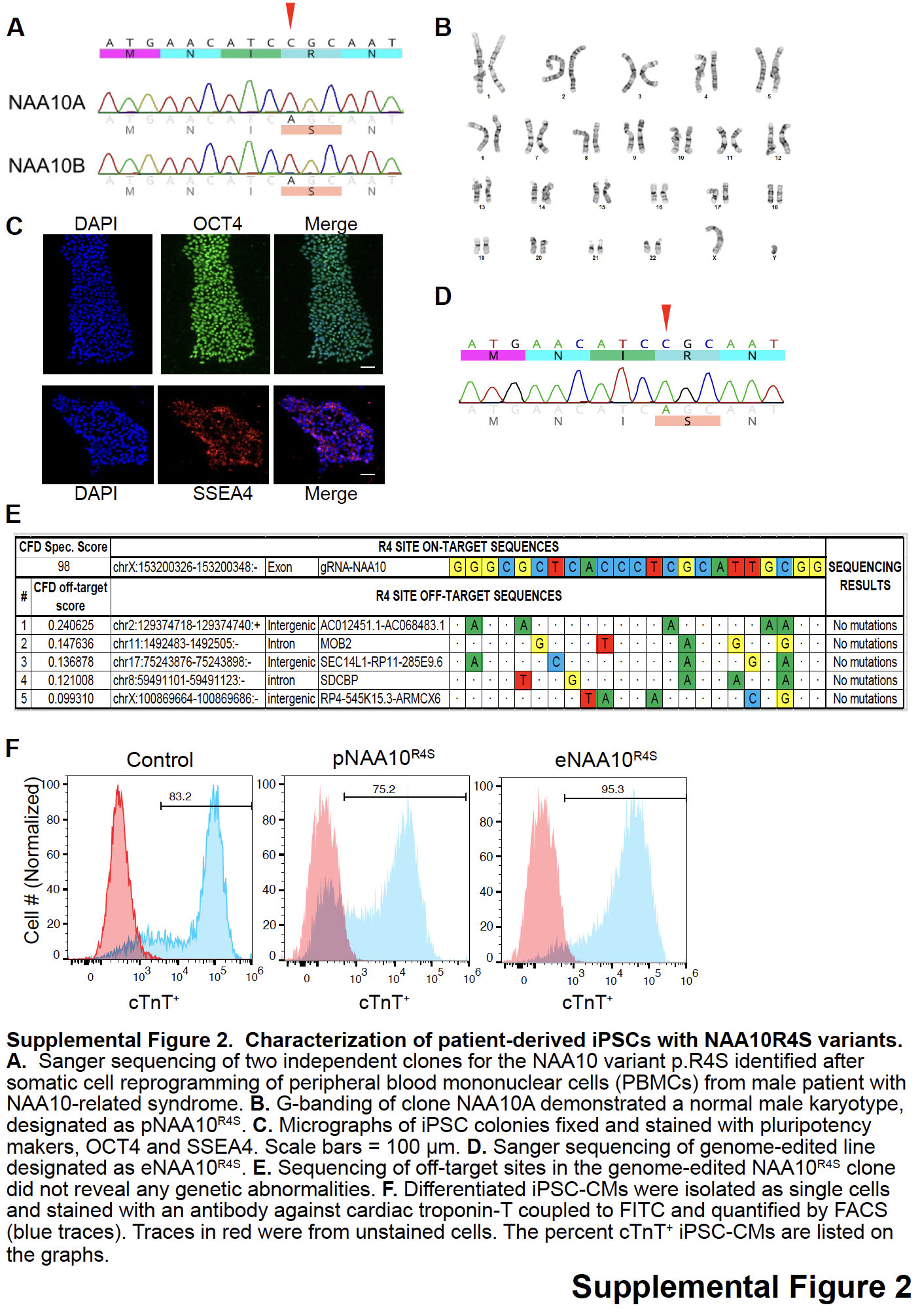

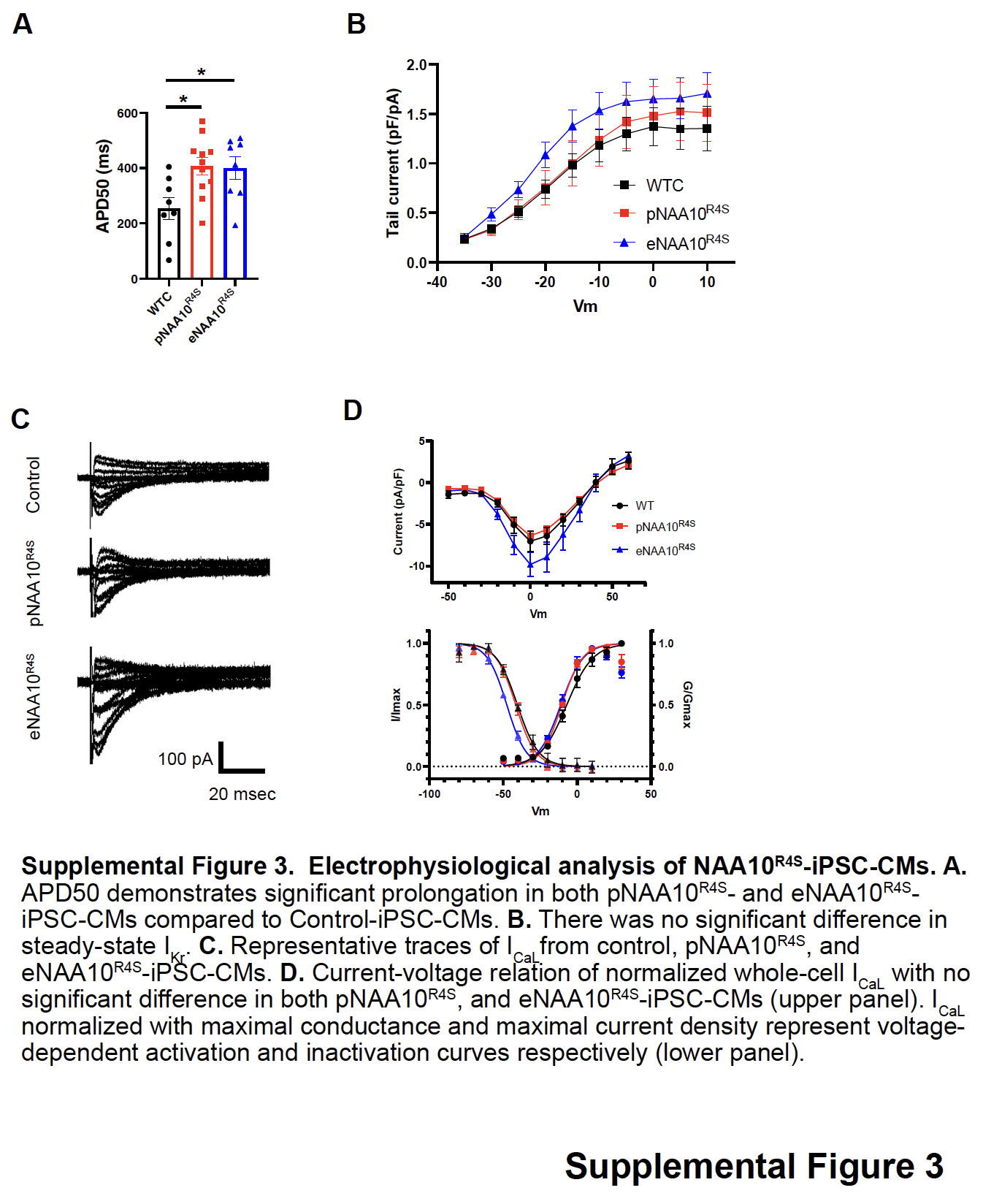

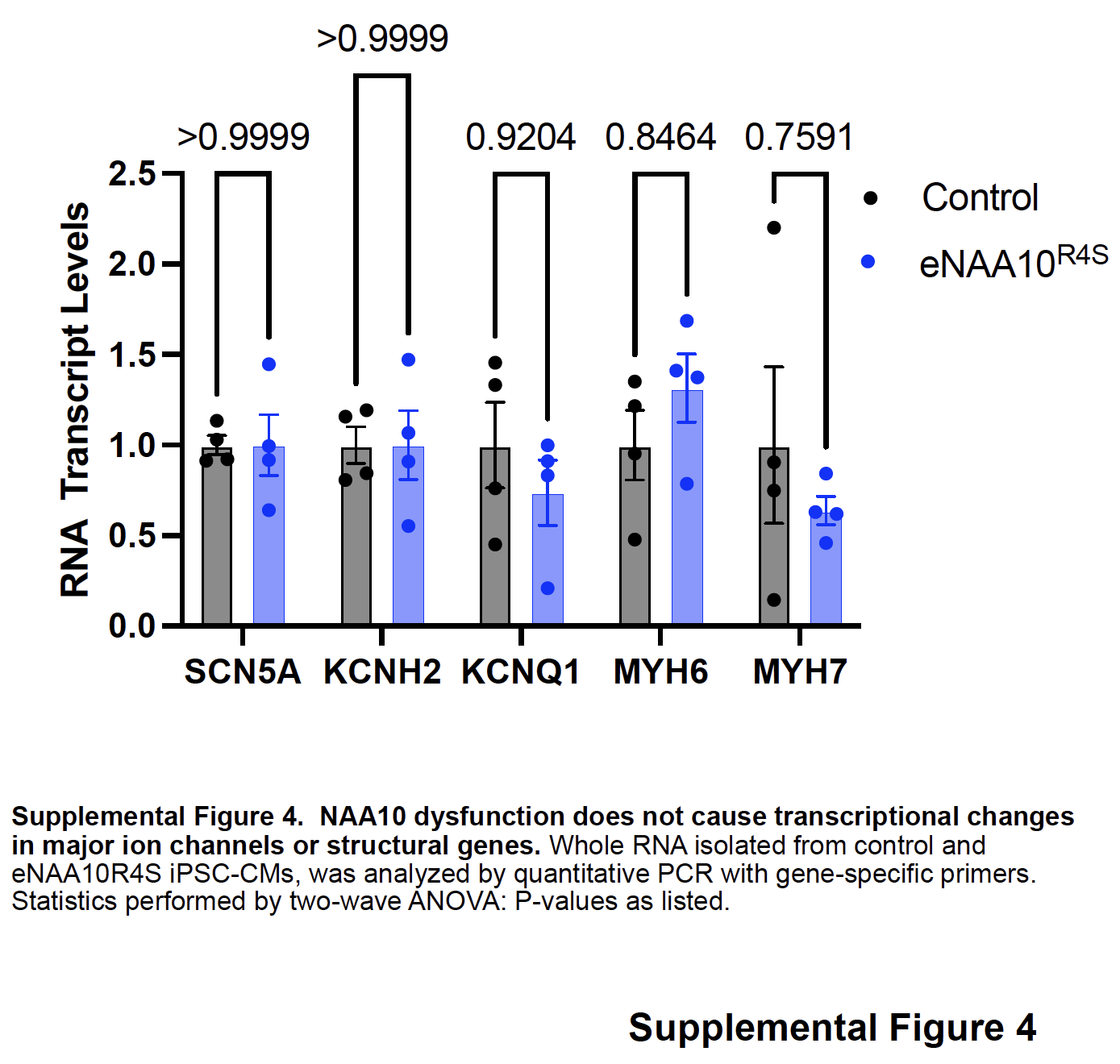

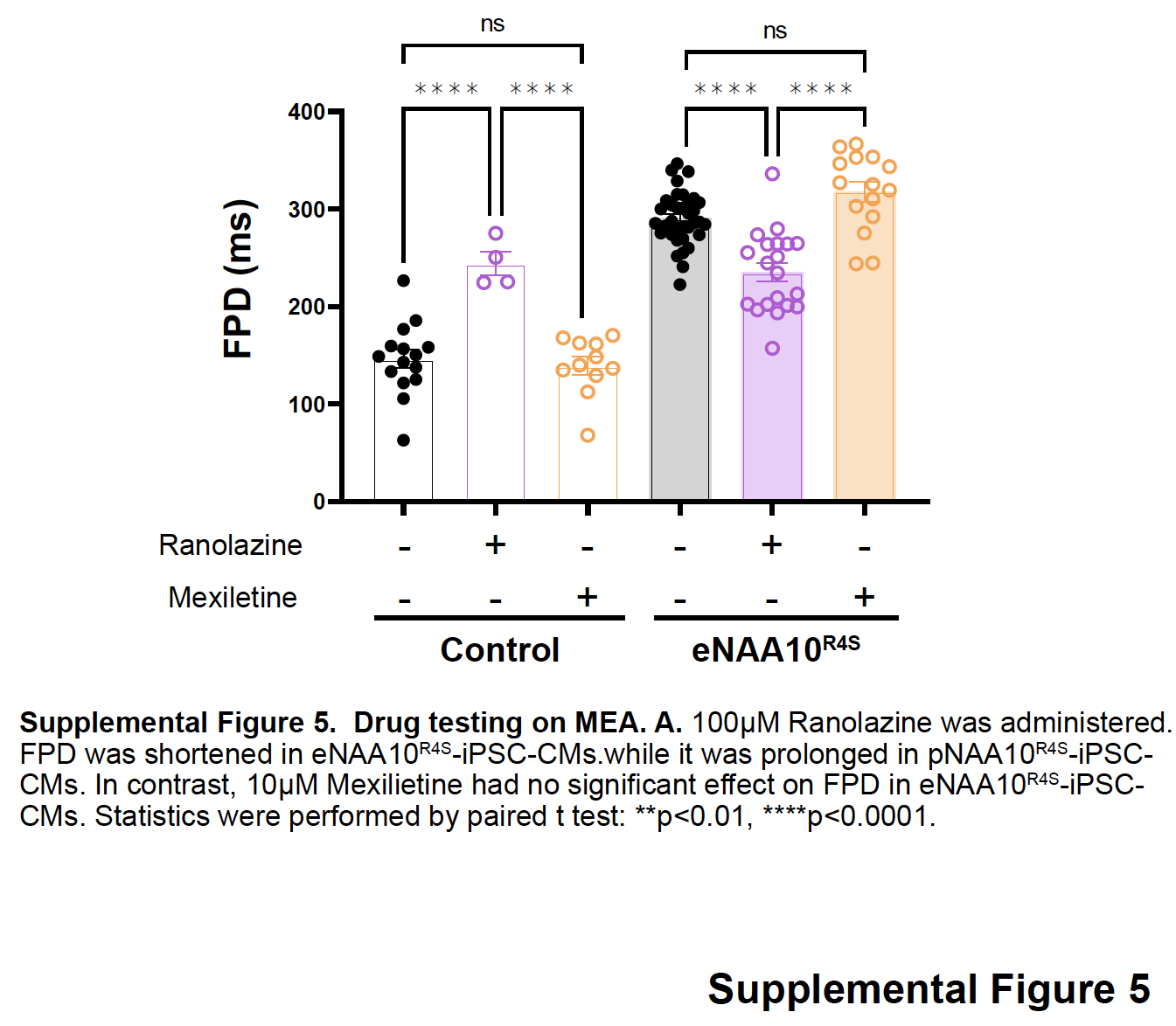

**SUPPLEMEMTAL REFERENCES**
